# In Step, the sole *Drosophila* TENT5 poly(A) polymerase, is required for spermatid elongation and individualization

**DOI:** 10.64898/2026.10.22.761032

**Authors:** Abdulqater Al-Nouman, Kyle Helms, Brook L. Falk, Julie A. Brill, Jennifer Curtiss

## Abstract

Cytoplasmic poly(A) tail length is a key regulator of mRNA translation and stability, controlled in part by non-canonical poly(A) polymerases of the TENT5 family, which are required for gametogenesis in mice. We characterize the sole *Drosophila melanogaster* TENT5 ortholog, which we name *in step* (*isep*). Although identified as a candidate target of the eye-specification factor Eyeless, *isep* mutants have normal eyes; instead, *isep* is essential for male fertility. The *isep* gene and its protein are expressed in spermatocytes and in elongating spermatids and are concentrated at the distal end of spermatid cysts. Mutant spermatids fail to elongate normally and to individualize. Whole-testis poly(A) tail sequencing revealed that, unexpectedly for the loss of a poly(A) polymerase, median tail length increased in mutants, with lengthened transcripts enriched for roles in axoneme assembly. Both wild-type and a predicted catalytically inactive Isep rescued sterility. We propose that Isep controls localized translation of axonemal transcripts at the spermatid tip, largely independently of its poly(A) polymerase activity.

**Article summary:** This work contributes to the *Drosophila* spermatogenesis community by identifying a previously unknown factor, *isep*, the fly’s only *TENT5*, which makes flies a model to study this enzyme family. This work traces the expression pattern of both *isep’s* transcripts and protein. Loss-of-function alleles generated by CRISPR-Cas9 mutagenesis result in complete male sterility, and immunofluorescence shows defects in spermatid elongation and individualization (separation into individual sperm). Mutant testes have longer median poly(A) tails, and lengthened transcripts are enriched for spermatogenesis and axonemal (sperm tail) genes. This work also suggests that the predicted poly(A) polymerase activity may be dispensable for spermatogenesis.

## INTRODUCTION

Most eukaryotic mRNAs carry a poly(A) tail, added co-transcriptionally in the nucleus by canonical poly(A) polymerases, and required for export to the cytoplasm. Once in the cytoplasm, the poly(A) tail can be lengthened or shortened, and these changes regulate gene expression post-transcriptionally by impacting translation efficiency and mRNA stability (Passmore and Coller, 2022). Accordingly, both deadenylases and non-canonical poly(A) polymerases (ncPAPs) exist in the cytoplasm to shorten and lengthen the poly(A) tail, respectively.

The best studied ncPAPs belong to the GLD-2/TENT2 family, which function in the germline and during early development in animals including *Caenorhabditis elegans*, *Xenopus laevis* and *Drosophila melanogaster* (Kadyk and Kimble, 1998, Wang et al., 2002, Nousch et al., 2017, Barnard et al., 2004, Rouhana et al., 2005, Benoit et al., 2008, Cui et al., 2008, Cui et al., 2013, Sartain et al., 2011, Liudkovska and Dziembowski, 2021). Research in *Xenopus laevis* places GLD-2 in a complex with the cytoplasmic polyadenylation element-binding protein (CPEB), where it lengthens the poly(A) tails of stored mRNAs required for oogenesis and early embryonic development, initiating their translation (Charlesworth et al., 2013, Ivshina et al., 2014).

However, the current picture appears more complicated. First, loss of the *D. melanogaster* GLD-2/TENT2 ortholog Wispy, which is clearly required for lengthening tails, does not globally reduce translational efficiency (Benoit et al., 2008, Cui et al., 2008, Cui et al., 2013, Eichhorn et al., 2016). Second, mouse GLD-2/TENT2 orthologs are dispensable for oocyte maturation, early embryonic development, and poly(A) tail elongation (Nakanishi et al., 2007). Instead, a different ncPAP family, TENT5, is important for oogenesis and spermatogenesis in mice (Brouze et al., 2024, Baptissart et al., 2026).

Like GLD-2/TENT2, TENT5s belong to the Terminal Nucleotidyl Transferase (TENT) superfamily (Liudkovska and Dziembowski, 2021). Vertebrate TENT5 members have poly(A) polymerase activity *in vitro* (Mroczek et al., 2017, Hu et al., 2020, Zhang et al., 2021), yet they are also required in processes independent of their poly(A) polymerase activity (Chen et al., 2020, Kazazian et al., 2020, Lai et al., 2024). Mammalian TENT5 orthologs can act as post-transcriptional regulators of secretory pathways, localizing to the ER and selectively polyadenylating transcripts encoding secreted, transmembrane, and ER resident proteins (Fucci et al., 2020, Viviani et al., 2026, Mroczek et al., 2017, Bilska et al., 2020, Herrero et al., 2020, Resnati et al., 2024, Gewartowska et al., 2021). In *C. elegans*, TENT-5 is expressed in male reproductive structures and polyadenylates male-specific transcripts, although its loss affects neither sperm morphology nor fertility (Mackiewicz et al., 2026). In mouse spermatogenesis, TENT5C localizes to the manchette, a transient microtubule structure between the spermatid head and tail, where it regulates transcripts encoding structural proteins (Zheng et al., 2019, Baptissart et al., 2026) In *D. melanogaster*, the two CPEB orthologs, Orb and Orb2, and one of the two GLD-2 orthologs, GLD-2, function during spermatogenesis (Sartain et al., 2011, Xu et al., 2012, Xu et al., 2014, Low et al., 2026). Because ncPAPs lack RNA-binding domains, they require RNA-binding proteins such as CPEBs for target specificity. Translationally repressed mRNAs are stored during the period of relative transcriptional quiescence in meiosis and spermatid morphogenesis (Kuhn et al., 1991, Schafer et al., 1993, Schafer et al., 1990, Schafer et al., 1995, White-Cooper, 2010, Raz et al., 2023). Thus, ncPAPs may be required to lengthen poly(A) tails of stored mRNAs during spermatogenesis, as they do during oogenesis.

Here we characterize the sole *D. melanogaster* TENT5 ortholog, which we named *in step* (*isep*). We show that *isep* is not required for eye development, where we initially identified it, but for spermatogenesis. *isep* transcripts and protein are present in spermatocytes and in spermatids, localizing to the distal ends of the spermatid cysts. Mutants for *isep* are male-sterile, with defects in spermatid elongation and individualization. Surprisingly, TAIL Iso-seq reveals that median poly(A) tail length is longer in *isep* mutants, rather than shorter, and that transcripts with significantly lengthened tails are enriched in axoneme assembly.

## RESULTS

### *isep* encodes a TENT5 non-canonical polyA polymerase

We identified *isep* (*CG46385*) while searching for transcriptional targets of the *Drosophila* Pax6 ortholog Eyeless (Ey). It was one of four genes shared between eye specific transcripts upregulated more than threefold by ectopic Ey expression in 3^rd^ instar wing imaginal disks (Nfonsam et al., 2012) and putative direct Ey targets identified by ChIP-seq in 3^rd^ instar larval eye-antennal imaginal disks (Yeung et al., 2018) (Fig. 1A); the other three, *eyes absent (eya), sine oculis (so),* and *distal antenna (dan)*, have established roles in eye development (Niimi et al., 1999, Bui et al., 2000, Zimmerman et al., 2000, Curtiss et al., 2007).

**Figure 1:**
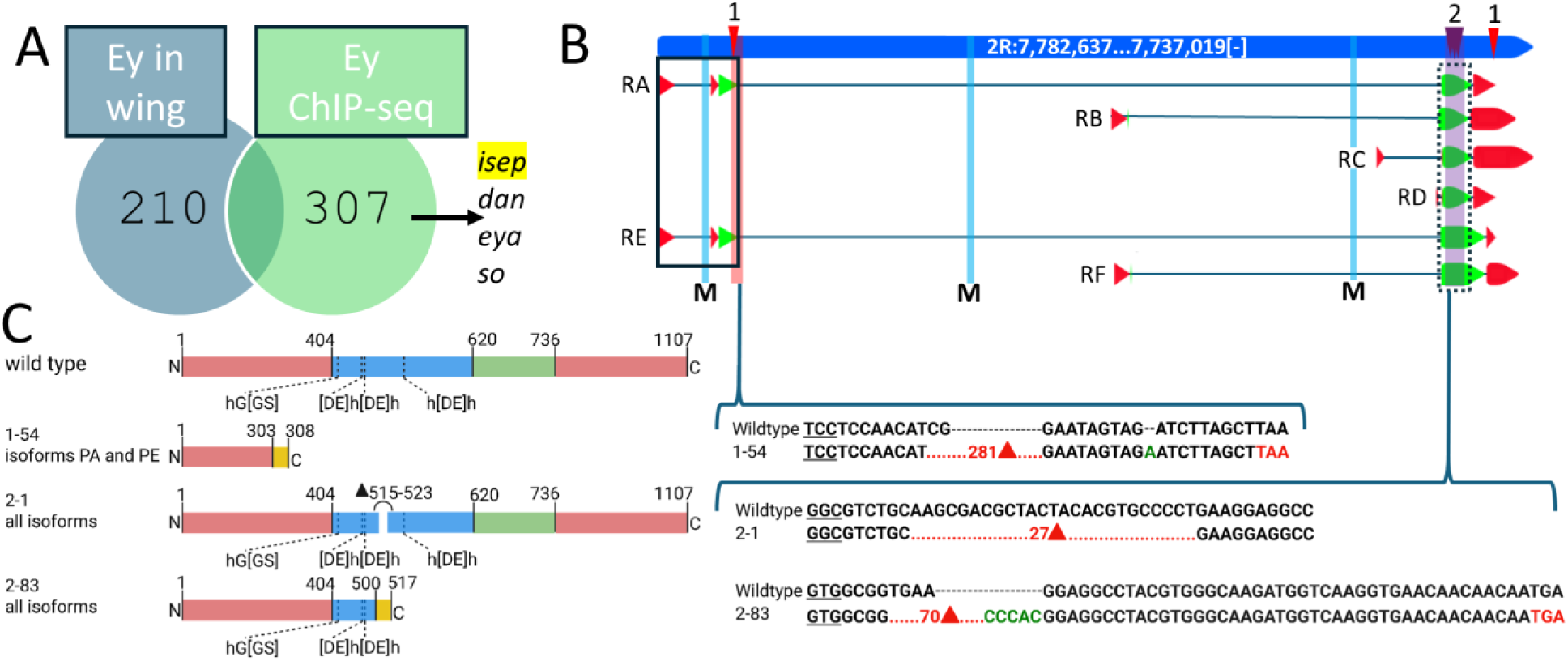
Identification of *isep*, CRISPR-Cas9 mutagenesis. **(A)** Venn diagram comparing eye-specific transcripts upregulated by ectopic Ey+Notch expression (left) and Ey ChIP-seq targets (right). **(B)** Genomic map of *isep* (blue) with the six mRNA isoforms below. Red wedges, non-coding exons; green wedges, coding exons; lines, introns. Red arrowheads, round 1 gRNAs; purple arrowheads, round 2 gRNAs; red and purple shading marks resulting deletions. Dashed box, catalytic domain shared by all isoforms; solid black box, predicted Ey binding region. Blue shading marked “M” indicates the three MIMIC insertions MI01603, MI03850, and MI12561 (left to right). Created in BioRender. Al-Nouman, A. (2026) BioRender.com/15l58lf. **(C)** Mutations corresponding to the shaded areas in **(B)**. DNA sequences are shown on the right: underlined codon indicates the reading frame; red triangles indicate the number of deleted base pairs; insertions are green; premature stop codons are red. Predicted protein products are shown on the left, with the N- and C-terminal extensions shown in red, NCD in blue, HD in green, and frameshifted residues in yellow. Dashed lines mark the five residues supporting Isep’s predicted poly(A) polymerase activity. Isoform PE was used for the protein models. Created in BioRender. Al-Nouman, A. (2026) k1wjmgz.

The 45.6 kb *isep* gene lies on chromosome 2R and encodes six mRNA isoforms producing five unique predicted proteins (Fig. 1B, FlyBase r6.52). Isoforms RC and RD differ in the 5′ and 3′UTRs but have identical coding sequences. RE encodes the largest predicted protein (1,107aa) and RC/RD the smallest (529aa). The RA/RE and RB/RF isoform pairs share the same transcription start site, exons, and translation start site but differ in their stop codons and 3′UTR, due to a predicted stop codon read-through identified by comparative genomics (Jungreis et al., 2016).

*isep* encodes a predicted non-canonical polyA polymerase (ncPAP) of the TENT5 family (Kuchta et al., 2016, Liudkovska and Dziembowski, 2021, Warkocki et al., 2018, Kuchta et al., 2009, Lacidogna et al., 2025) (Fig. S1). Vertebrates possess four orthologs (TENT5A-D), whereas *D. melanogaster* has only one. Like other TENT5 family members, Isep contains an N-terminal catalytic domain (NCD) with three conserved nucleotidyltransferase motifs (hG[GS], [DE]h[DE]h, and [DE]h) and a C-terminal helical domain (HD) (Fig. 1C), but lacks the RNA-binding domain (RBD) found in canonical PAPs (Liudkovska and Dziembowski, 2021). Using the crystal structure of the *Xenopus tropicalis* TENT5B ortholog as a model (Hu et al., 2020), Isep retains all five residues required for TENT5B’s poly(A) polymerase activity and seven of the eight positively charged residues implicated in RNA binding (Fig. S1 and legend). Thus, Isep likely functions as a non-canonical polyA polymerase.

Isep also has extended N- and C-terminal regions (Fig. 1C, Fig. S1) predicted by the ParSe 2.0 tool (Wilson et al., 2023) to be intrinsically disordered, with portions predicted to drive phase separation (Fig. S1). The first disordered domain after the HD is glutamine-rich, and the PANTHER database (Thomas et al., 2022) classifies Isep as a prion-like protein with a Q/N-rich domain. Together, these features suggest Isep can undergo phase separation (Gomes and Shorter, 2019, Ibrahim et al., 2023, Chong and Mir, 2021).

### *isep* is required for male fertility

To determine *isep* gene function, we used a CRISPR/Cas9-based strategy (Gratz et al., 2015b, Gratz et al., 2015a) to generate three *isep* mutant alleles (Fig. 1C, see Methods). *isep^1-54^* deletes both the predicted NCD and HD but is expected to affect only the RA/RE isoforms (Fig. 1B,C). *isep^2-1^*removes 9 amino acids from the predicted NCD of all six isoforms (Fig. 1C) but leaves all poly(A) polymerase catalytic residues and putative RNA-binding residues intact. *isep^2-83^* (Fig. 1B) truncates approximately the C-terminal half of the NCD and the entire HD, including the h[DE]h catalytic motif and all the putative RNA-binding residues (Fig. 1C). All three alleles are viable *in trans* to Df(2R)ED1725 and Df(2R)Exel6054 (Ryder et al., 2007, Parks et al., 2004) and to three different MiMIC insertion mutants (Fig. 1B) (Nagarkar-Jaiswal et al., 2015, Venken et al., 2011), with no eye or other external defects. However, the *isep^2-83^*allele causes male infertility.

Fertility was tested by crossing males and females of each *isep* allele to *w^1118^* flies of the opposite sex, with *w^1118^*males and females crossed to each other as a control. Control crosses resulted in an average hatch rate of 94% (Fig. 2A,A’). In contrast, *w^1118^* females mated to *isep^2-83^* homozygous males had a 0% hatch rate (Fig. 2A,A’), whereas *isep^2-83^* homozygous females crossed to *w^1118^* males had an average hatch rate of 97% (Fig. 2A,A’). Thus, *isep^2-83^* homozygous males were sterile while homozygous females were fertile. Male and female *isep^1-54^* and *isep^2-1^*homozygotes were fertile, consistent with *isep^1-54^* affecting only 2 of 6 isoforms and *isep^2-1^* deleting a small number of non-catalytic residues.

**Figure 2:**
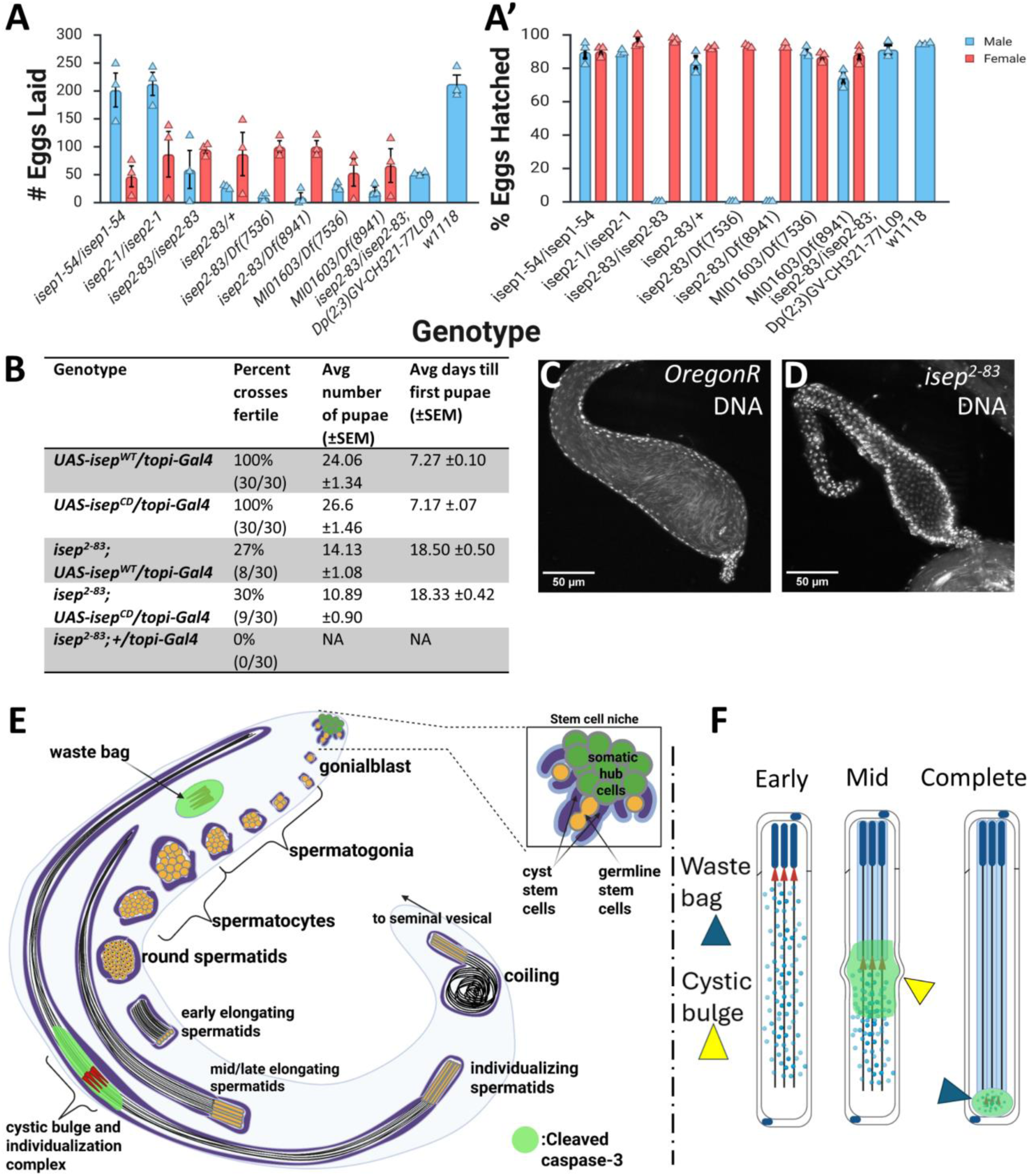
*isep* is required for male fertility. Egg lay assays showing **(A)** the average number of eggs laid and **(A’)** the percentage hatched for each genotype, with males in blue and females in red. Triangles are replicates; error bars, SEM. **(B)** Rescue assay showing the percentage of individual male crosses rescued for each genotype, the average number of pupae, and days to emergence of the first pupa. **(C,D)** Hoechst staining of seminal vesicles from 3-5-day-old males; wild-type mature sperm are characterized by long, thin nuclei. **(E)** Stages of spermatogenesis from stem cell divisions at the hub to individualization. Created in BioRender. Al-Nouman, A. (2026) wiejcjv. (**F)** Simplified model of individualization. Created in BioRender. Al-Nouman, A. (2026) BioRender.com/3h5140l.

To further assess the effect of *isep^2-83^* on fertility, we tested heterozygotes and *isep^2-83^* males *in trans* over each deficiency. Heterozygous males were fertile with a 97% hatch rate, whereas *isep^2-83^*/Df(2R)ED1725 or *isep^2-83^*/Df(2R)Exel605 males yielded a 0% hatch rate (Fig. 2A,A’), establishing *isep^2-83^*as a recessive loss-of-function allele. Fertility was restored by an interchromosomal duplication spanning cytological bands 43E13-43E18 that contains *isep* (Fig. 2A,A’).

Three MiMIC gene trap alleles (Nagarkar-Jaiswal et al., 2015) did not cause male sterility *in trans* to either deficiency (not shown and Fig. 2A,A’). *MI12561* is not predicted to truncate any isoform, as it lies between two non-coding exons at the 5’ end of the RA and RE isoforms.

*MI03850* is predicted to truncate only the RA and RE products. In contrast, *MI01603* is predicted to truncate all isoforms except those encoded by RC and RD. The *isep^2-83^*allele is predicted to affect all isoforms and causes male sterility, while alleles sparing only RC and RD do not, suggesting that RC/RD are sufficient for normal spermatogenesis in *Drosophila*. Consistent with this, Fly Atlas 2 (Leader et al., 2018) shows that all isoforms are expressed in the testis, with RD as the only isoform exclusively expressed there.

To assess whether *isep^2-83^* sterility reflects a spermatogenesis defect, we labeled seminal vesicles from 3-5 day-old males with Hoechst to mark nuclei. Whereas the needle-like nuclei of mature sperm were readily apparent in wild-type, *isep^2-83^* homozygous seminal vesicles appeared deflated and empty with no detectable sperm nuclei (Fig. 2C,D).

To test whether Isep’s poly(A) polymerase catalytic activity is required, we generated two UAS constructs encoding wild-type Isep or a predicted catalytically dead version (see Methods), as substitution of one of the three catalytic residues abolishes PAP activity (Zhang et al., 2021).

Both constructs were driven with *topi-Gal4*, which is active in primary spermatocytes (Galletta et al., 2020). Each construct rescued *isep^2-83^* sterility in approximately one-third of crosses, with a considerable lag (∼18 days vs ∼7 days for controls) in the appearance of the first pupa and fewer pupae overall (Fig. 2B), suggesting a partial rescue that we attribute to Gal4/UAS timing or a delay in spermatogenesis. However, the wild-type and catalytic-dead constructs behaved comparably, suggesting that Isep’s poly(A) polymerase activity is dispensable or secondary to a structural role.

### *isep* is expressed in spermatocytes and post-meiotically in elongating spermatids

At the tip of the *D. melanogaster* testis, somatic hub cells contact germline and cyst stem cells (Fig. 2E). Each displaced germ cell becomes a gonialblast that undergoes four rounds of mitosis and then meiosis to produce 64 round spermatids (Fuller, 1993). Round spermatids elongate their flagellar axonemes and remodel and compact their nuclei through the deposition of protamines and other non-histone proteins (Fabian and Brill, 2012, Fuller, 1993).

Because germ cell cytokinesis is incomplete, spermatids remain syncytial until individualization, when actin cones assemble around all 64 nuclei and form an individualization complex that travels down the length of each spermatid within a cystic bulge, stripping away unneeded organelles and cytoplasm into a waste bag and encasing each spermatid in its own membrane (Fig. 2E,F) (Tokuyasu et al., 1972, Fabrizio et al., 1998, Fuller, 1993, Fabian and Brill, 2012). A non-apoptotic caspase cascade is also essential, with activity restricted to the cystic bulges, the regions distal to them, and the waste bags (Arama et al., 2003, Arama et al., 2006). Although most transcription halts before the meiotic divisions, the cup and comet genes are post-meiotically transcribed in mid-elongation cysts and localize to the distal ends of elongating spermatids (Barreau et al., 2008, Lim et al., 2012, Olivieri and Olivieri, 1965, Gould-Somero and Holland, 1974, Raz et al., 2023).

We used HCR-FISH probes targeting a region common to all isoforms and co-stained with Hoechst to determine where *isep* is expressed in the testis (Fig. 3). *isep* transcripts were absent from the hub and spermatogonia (Fig. 3A, white arrowhead) but present in primary spermatocytes and persisted through the meiotic divisions (Fig. 3B-C, white and magenta arrows). Transcripts were also detected in elongating spermatids, with the strongest signal at the distal end of the cyst and tapering more proximally in a comet-like expression pattern (Fig. 3A-C, yellow arrows). In cysts further along in development, during or at the end of elongation, localization shifted to a cup-like pattern concentrated at the distal end (Fig. 3A, yellow arrowheads). Thus, *isep* is expressed in primary spermatocytes and post-meiotic spermatids, like other cup and comet genes.

**Figure 3:**
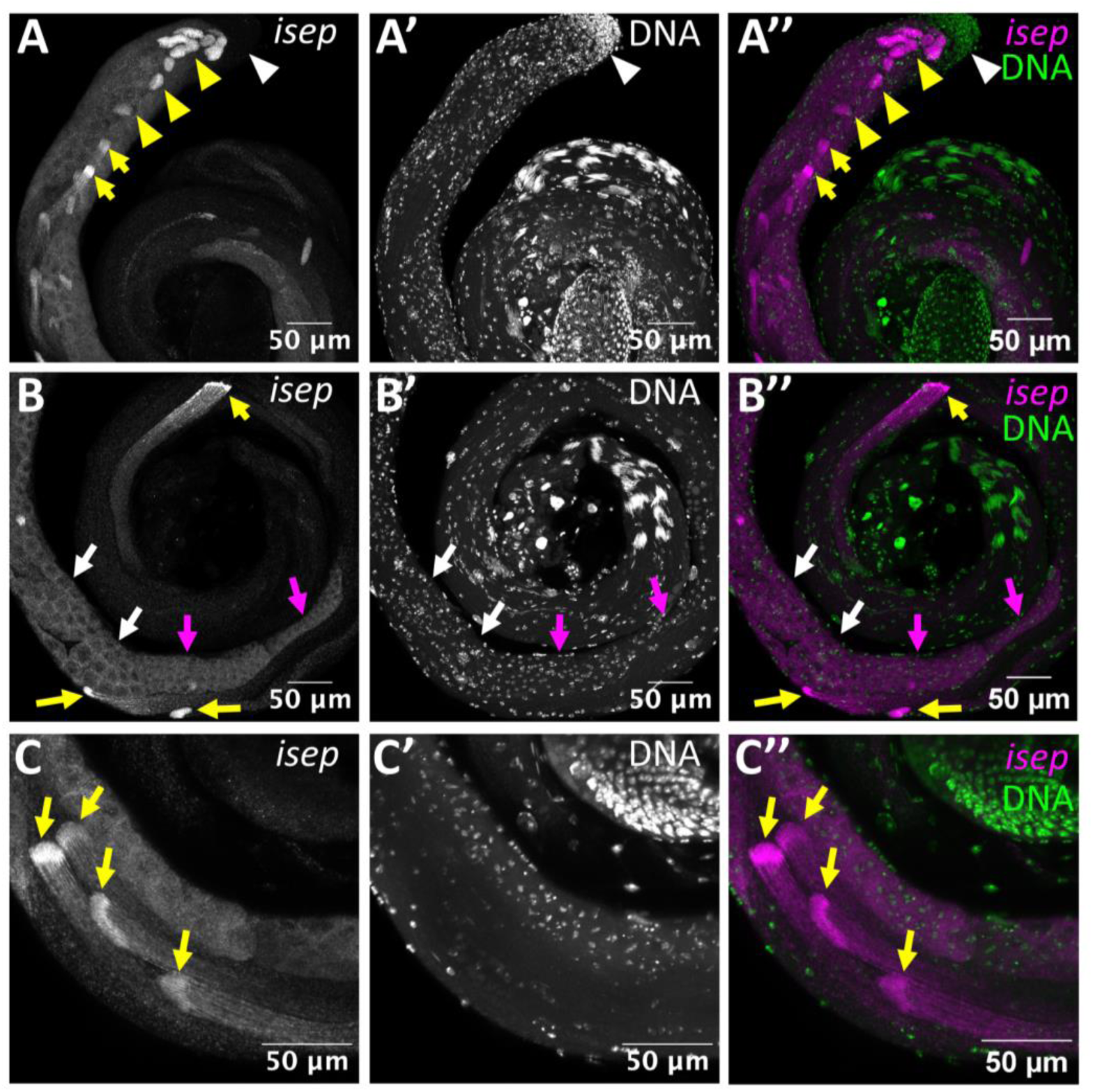
HCR-FISH in the testis probed for *isep* (magenta) co-stained with DNA (Hoechst, green). **(A)** White arrowheads, testis tip (hub) and mitotically dividing cells. *isep* transcripts at the distal end of cysts, showing a comet-like pattern (yellow arrows) and a cup-like pattern where the distal ends of the cysts are near the hub (yellow arrowheads). **(B)** White arrows, primary spermatocytes; magenta arrows, spermatocytes that have initiated meiosis; *isep* transcripts at the distal end of a cyst, showing a comet-like pattern (yellow arrow). **(C)** *isep* transcripts at the distal ends of spermatid cysts, showing a comet-like pattern (yellow arrows).

### Isep protein is expressed in spermatocytes and in elongating spermatids

To visualize when and where Isep protein is expressed, we endogenously tagged Isep with mScarlet3 using the SEED/Harvest method (Aguilar et al., 2024) (see Methods). Homozygous tagged males were fertile and displayed no apparent spermatogenesis defects, indicating that the internal fluorescent tag does not compromise Isep function. We detected the protein at the same stages as its transcripts. We co-stained with an antibody against Orb2, the *Drosophila* CPEB ortholog that is present in primary spermatocytes and elongating spermatid cysts (Xu et al., 2012). Isep and Orb2 were co-expressed in the cytoplasm of primary spermatocytes (Fig. 4A,B), but Isep declined in the cytoplasm of cells that had completed meiosis, whereas Orb2 persisted (Fig. 4A,B, red dashed outlines). Additionally, in primary spermatocytes, Isep exhibited a striking nuclear localization pattern (Fig. 4C) in the region where the Y chromosome is actively transcribed (Ball et al., 2023, Bonaccorsi et al., 1988).

**Figure 4:**
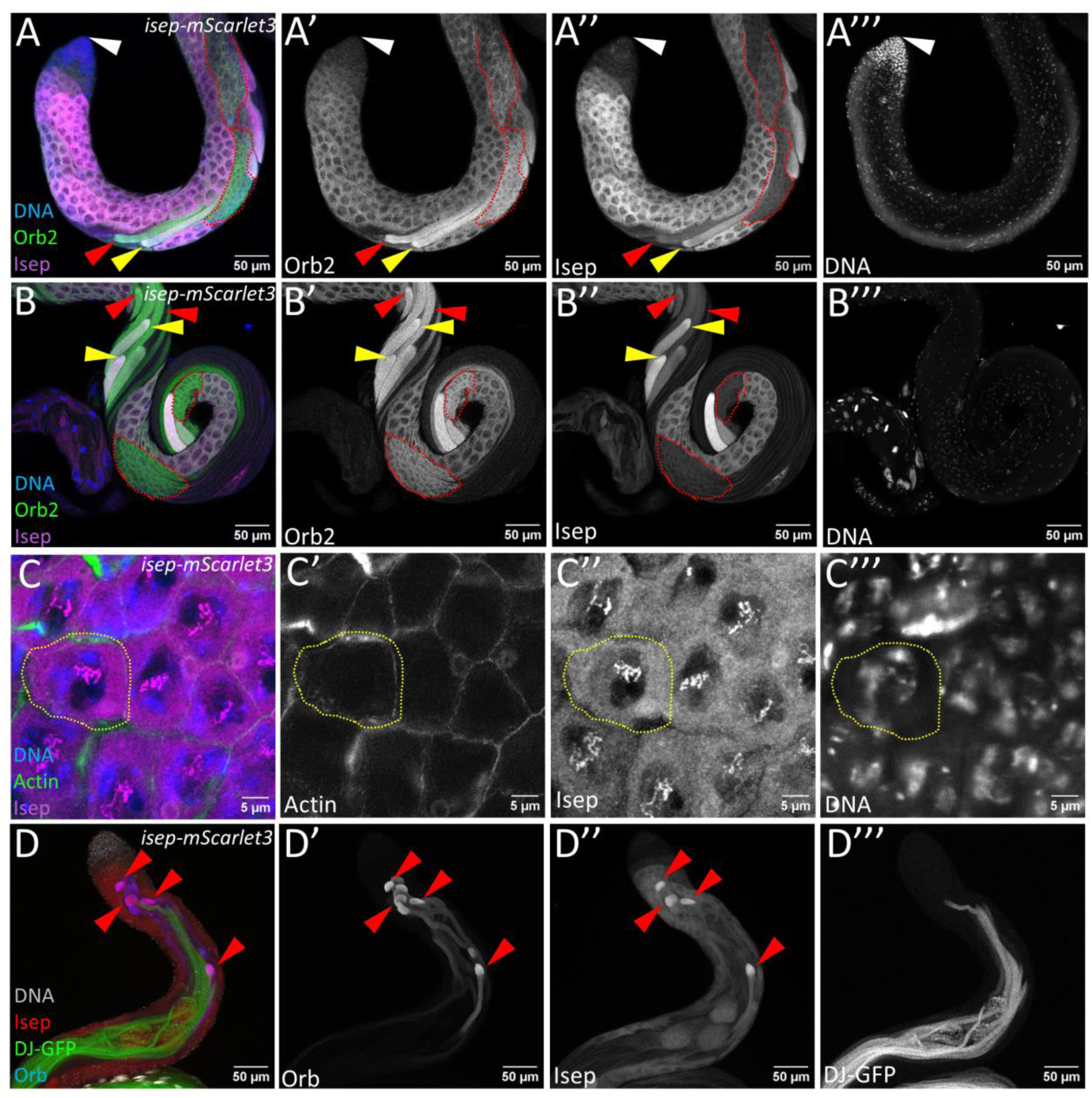
Isep protein localizes in the cytoplasm and nucleus of primary spermatocytes and at the distal end of elongating spermatid cysts. **(A,B)** Testis regions showing Isep (magenta) and Orb2 (green). White arrowhead, testis tip; red dashed outlines, cysts with higher Orb2 than Isep; red arrowheads, elongating cysts with higher Orb2 relative to Isep; yellow arrowheads, elongating cysts with comparable levels of Isep and Orb2. **(C)** Primary spermatocytes co-stained for Isep (magenta), actin (phalloidin, green), and DNA (Hoechst, blue); yellow dashed circle outlines a single spermatocyte**. (D)** Isep (red) at the distal ends of late elongating cysts before individualization, as indicated by the absence of DJ-GFP (green); red arrowheads indicate cysts with both Orb (blue) and Isep.

After meiosis, Isep and Orb2 were both readily detected starting at mid-elongation and co-localized along the length of the cysts (Fig. 4A,B, yellow arrowheads). As elongation progressed, Isep decreased relative to Orb2 (Fig. 4A,B, red arrowheads). Because *isep* transcripts were still present at this stage (Fig. 3B, magenta arrows), the decrease in protein may reflect post-transcriptional regulation. After elongation, a delay occurs between the disappearance of Orb2 and the onset of individualization markers such as Don Juan (DJ) (Santel et al., 1998), and this interval is marked by Orb, the other *Drosophila* CPEB ortholog (Xu et al., 2012). Isep was detected again in some cysts during this window (Fig. 4D, red arrowheads), colocalizing with Orb mainly at the distal end of the cysts and forming a gradient towards the proximal end. In summary, Isep first appears in primary spermatocytes, decreases after meiotic divisions, reappears with Orb2 during early elongation, and after a further drop rises again with Orb.

### Loss of *isep* function results in individualization failure

Because Isep is initially expressed in primary spermatocytes, we asked whether meiosis proceeds normally in *isep^2-83^* mutants. Phase contrast imaging of live squashes revealed round spermatids with no apparent morphological defects and the correct 1:1 ratio of nuclei to nebenkerne (Fig. 5A,B), suggesting meiotic divisions are unaffected.

**Figure 5:**
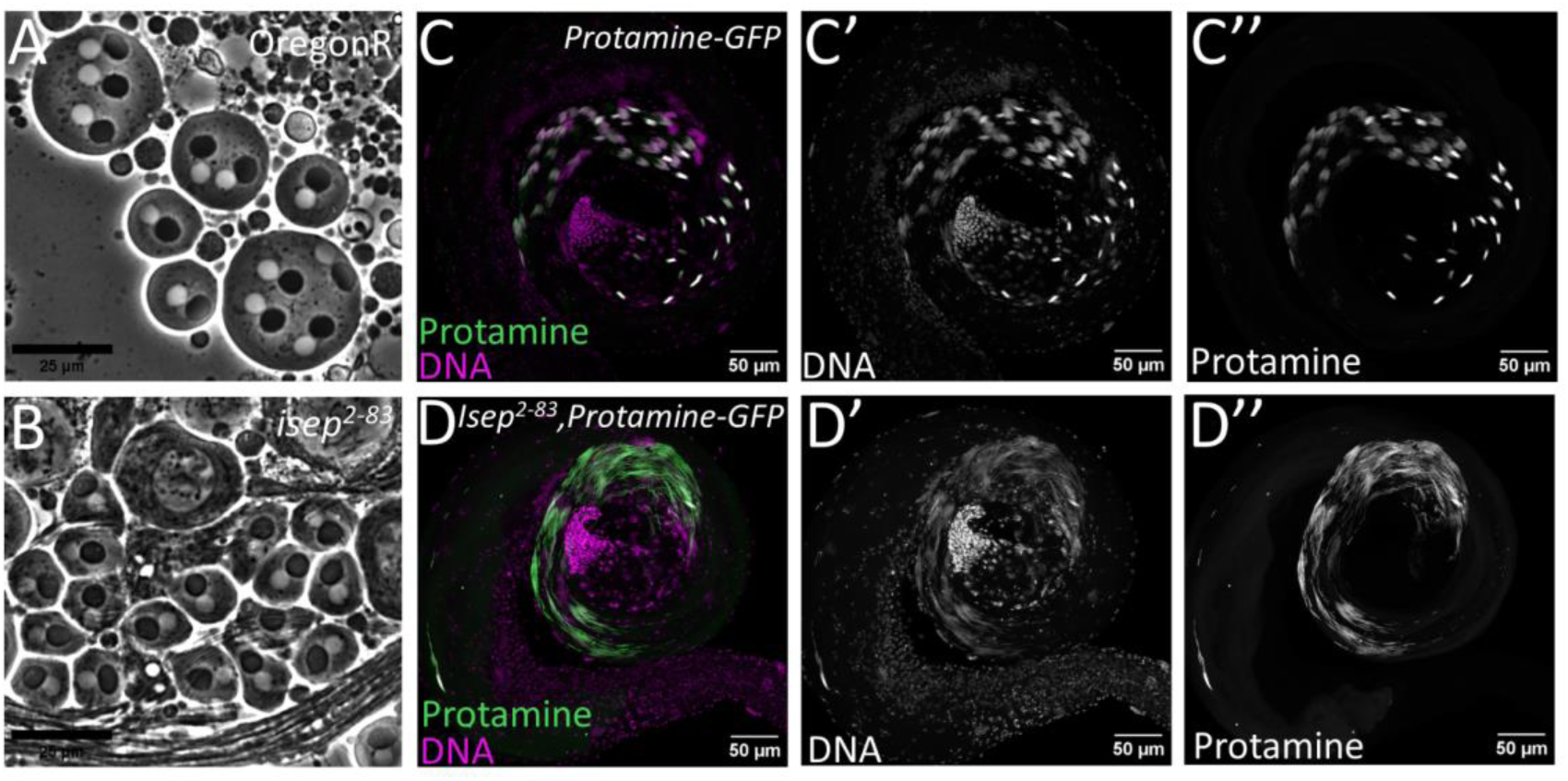
*isep* loss affects neither meiosis nor protamine incorporation. **(A,B)** Phase-contrast images of live squashed preparations of wild-type **(A)** and *isep^2-83^***(B)** testes showing round spermatids with the appropriate (1:1) nebenkern (dark) to nucleus (white) ratio. **(C-C’’,D-D’’)**. Testes expressing eGFP-tagged protamine B (green) and co-stained for DNA (Hoechst, magenta); nuclei are scattered in *isep^2-83^*, although protamine B incorporation is not affected.

Loss of TENT5C causes headless sperm in mice (Zheng et al., 2019), and TENT5C associates with the centrosome regulator Plk4 to inhibit its activity, independent of its poly(A) polymerase activity (Chen et al., 2020, Kazazian et al., 2020). Because centrosome-derived basal bodies link the axoneme to the nucleus (Galletta et al., 2020, Fabian and Brill, 2012), defects in centrosome dynamics can result in headless sperm, a phenotype reported for knockdown of the ncPAP GLD2 in testes (Sartain et al., 2011). Using the basal body marker *asterless-GFP (asl-GFP)* (Galletta et al., 2016), we observe no docking defects; basal bodies remained closely associated with canoe-stage spermatid nuclei in wild-type and *isep^2-83^* (Fig. S2). Furthermore, Protamine B-GFP was also normally incorporated into *isep^2-83^*nuclei (Fig. 5C,D), suggesting that nuclear compaction occurs normally. Thus, meiosis, basal body docking, nuclear shaping, and compaction all occur normally in *isep^2-83^* mutants.

Strikingly, however, whereas protamine-labeled nuclei in wild-type late-stage cysts are tightly packed, nuclei in *isep^2-83^*are scattered and appear highly disorganized (Fig. 5C,D). This phenotype resembles mutants that fail to individualize. We analyzed individualization directly by staining with an antibody against the active, cleaved form of caspase-3. In wild-type cysts, caspase activity was restricted to the cystic bulge, the region distal to the cystic bulge (Fig. 6A’, white dashed boxes), and waste bags (Fig. 6A’, yellow dashed outlines). In *isep^2-83^*, caspase-3 activity was uniform throughout the cyst, the cystic bulges appeared elongated and flattened, and waste bags were absent (Fig. 6B’). Phalloidin staining showed that actin cones in wild-type cysts were tightly organized in individualization complexes (Fig. 6A’’,A’’’) but scattered in *isep^2-83^* and failed to travel as a single unit (Fig. 6B’’,B’’’), although their morphology appeared normal. Scattered actin cones, absent waste bags, and uniform caspase-3 activity together indicate a failure of spermatid individualization; the scattered actin cones inspired the name *in step* (*isep)*.

**Figure 6:**
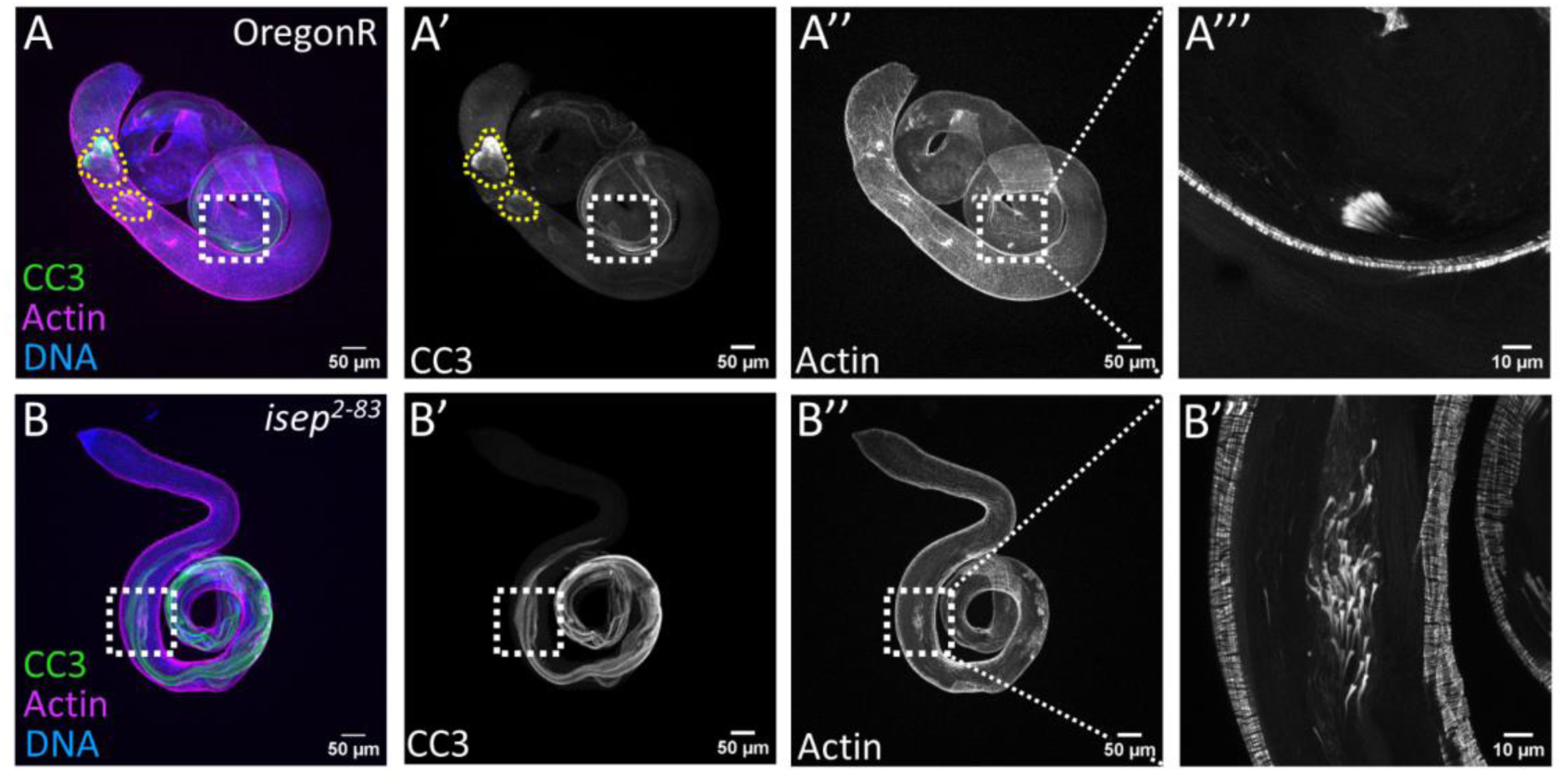
*isep^2-83^* mutant spermatids fail to individualize. **(A,B)** Wild-type and *isep^2-83^* testes stained for cleaved caspase 3 (CC3, green), actin (phalloidin magenta), and DNA (Hoechst, blue). Insets show the individualization complex in greater detail. **(A-A’’’)** In wild type, caspase activity is restricted to the cystic bulge and distal regions (white dashed box); actin cones travel synchronously; and waste bags form (yellow dashed circles). **(B-B’’’)** *isep^2-83^* mutant testes exhibit ubiquitous cleaved caspase 3 throughout the cysts, scattered actin cones, flatter cystic bulges (white dashed box), and an absence of waste bags.

Next, we asked whether individualization begins normally by tracing early, mid, and late stages using Hoechst, phalloidin, and anti-polyglycylated tubulin antibodies, which mark axonemes during and after individualization (Bressac et al., 1995, Bre et al., 1996). In wild-type controls, actin cones assembled neatly at the base of each spermatid nucleus in the basal region of the testis, then traveled synchronously, reaching the midpoint of the cyst and finally the tip of the testis (Fig. 7Ai-iii,A’i-iii). In *isep^2-83^*, nuclei and actin cones were already scattered at the earliest stage and became progressively more so as the cones traveled towards the testis tip (Fig. 7Bi-iii,B’i-iii). This phenotype resembled that of *orb2* mutants lacking the zinc finger domain (*orb2^ΔZZ^*) (Low et al., 2026), in which scattering likewise begins while actin cones are still proximal to the nuclei, and scattering spans a larger portion of the testis than *soti* or *eIF4E-*5 mutants (Kaplan et al., 2010, Shao et al., 2023).

**Figure 7:**
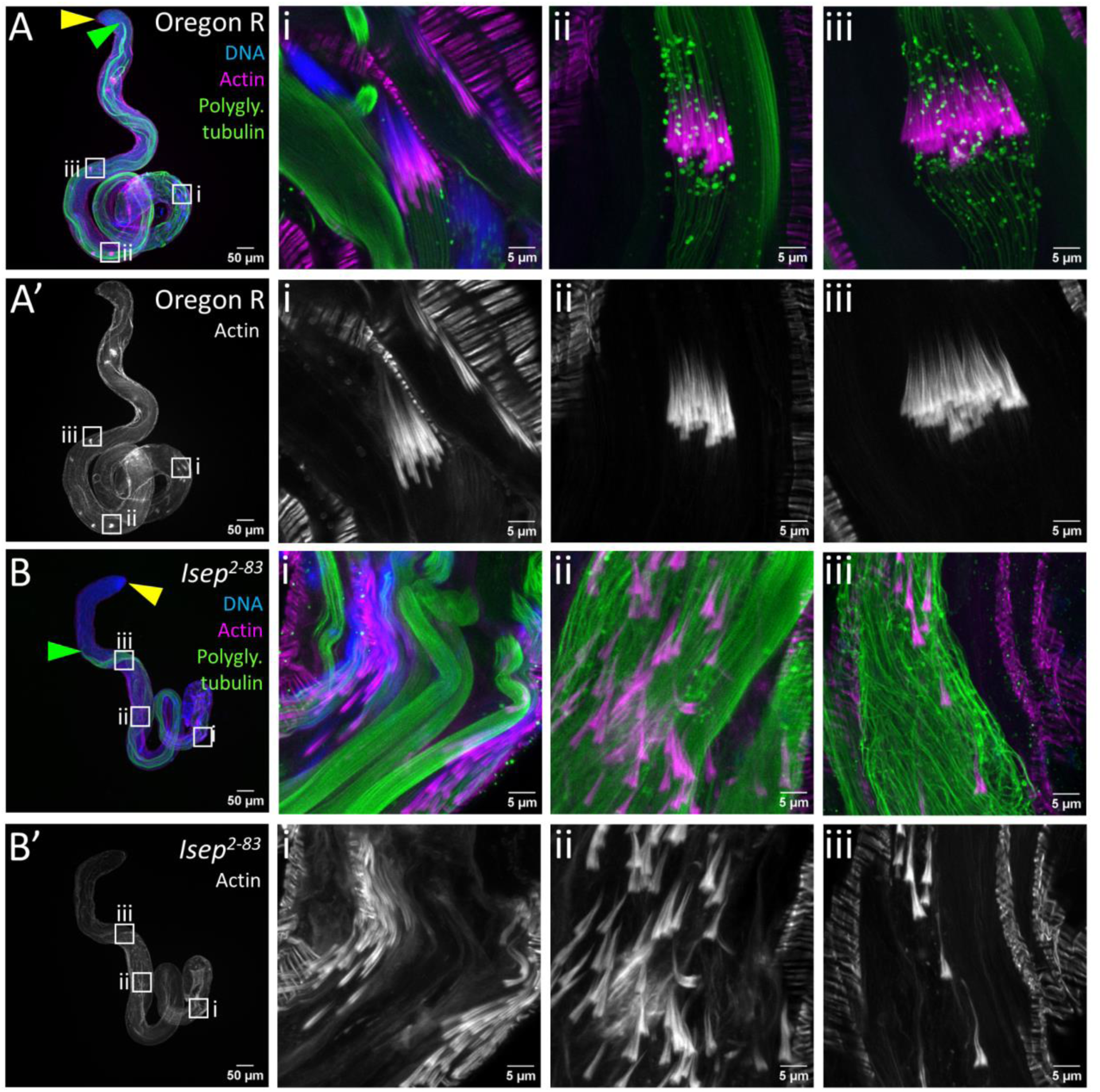
*isep^2-83^* mutant spermatids display actin and nuclear scattering at the onset of individualization. **(A-B)** Testes stained for polyglycylated tubulin (green), F-actin (phalloidin; magenta), and DNA (Hoechst; blue). Yellow arrowheads mark the testis tip. Green arrowheads mark the first polyglycylated tubulin signal relative to the testis tip. Insets: Early **(i),** middle **(ii),** and late **(iii)** stages of individualization, rotated to a common orientation. **(A’i)** Wild-type actin cones form around the base of spermatid nuclei and travel synchronously **(A’ii-iii). (B’i)** *isep^2-83^* actin cones are scattered from the earliest stage and become progressively more so **(B’ii-iii)**. **(Biii)** Microtubules at the distal ends of the cysts appear broken and disorganized.

### *Isep^2-83^* cysts show defects before individualization

Given that defects appeared at the onset of individualization, we examined Soti, a regulator of caspase activation. Localization of *soti* transcripts in a comet pattern generates a gradient of Soti protein highest at the distal end of the cyst, which helps establish an opposing gradient of active caspase. Loss of *soti* causes individualization failure with ubiquitous caspase activity and scattered actin cones (Kaplan et al., 2010, Barreau et al., 2008), similar to what we observed in *isep^2-83^*. Both eIF4E-5 and Orb2 are also required for individualization and proper Soti accumulation at the distal end of the cyst (Low et al., 2025, Shao et al., 2023). Anti-Soti staining revealed a similar defect in *isep^2-83^*; Soti failed to accumulate at the distal end of the cyst (Fig. 8A,B, magenta arrowhead). The Soti pattern also showed that elongating *isep^2-83^* cysts did not extend to the tip of the testes, as in wild type (Fig. 8A’,B’, magenta and yellow arrowheads).

**Figure 8:**
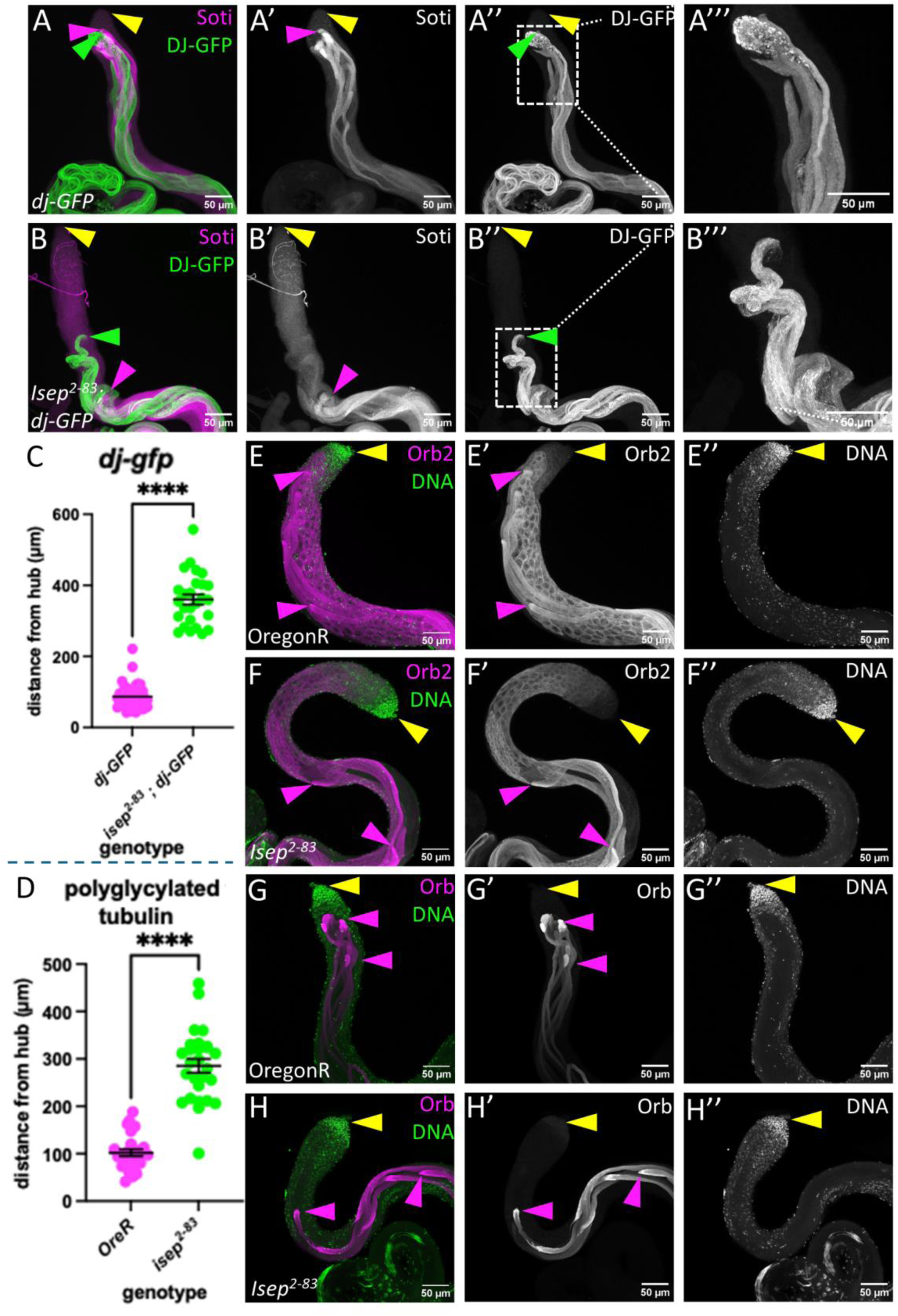
Proteins enriched at the distal ends of cysts are mislocalized in *isep^2-83^* mutants, and axonemal markers do not extend as far back towards the hub. **(A-B)** Wild-type **(A)** and *isep^2-83^*(B) testes in a DJ-GFP background (green) stained for Soti (magenta). Yellow arrowhead, testis tip; green arrowhead, first DJ-GFP signal relative to the testis tip; magenta arrowhead, first Soti signal relative to the testis tip. **(A’, B’)** Soti in pre-individualizing cysts, forming a gradient highest at the distal ends of cysts. **(A’’, B’’)** DJ-GFP marking individualizing to post-individualized spermatids; **(A’’’, B’’’)** insets highlighting the disorganized/broken tail phenotype. **(C-D)** Quantification of the distance from the hub to the first DJ-GFP **(C)** or polyglycylated tubulin **(D)** signal for each genotype; **** p < 0.0001. **(E-F)** Testes stained for Orb2 (magenta), which localizes to the distal ends along the cysts as well as spermatocytes, and DNA (Hoechst, green). Yellow arrowheads, testis tips; magenta arrowheads, first and last Orb2 in elongating cysts. **(G-H)** Testes stained for Orb (magenta) and DNA (Hoechst, green), with the same arrowhead markings as in (E-F); Orb forms a gradient similar to Soti in wild type **(G)** but in *isep^2-83^* does not extend as far back and adopts a flared paintbrush morphology **(H)**.

Because Soti diminishes as individualization begins, we further examined fully elongated cysts expressing Don Juan-GFP (DJ-GFP), which localizes to mitochondrial derivatives along the axoneme, from individualization onwards (Santel et al., 1998). DJ-GFP confirmed that axonemes did not extend nearly to the testis tip, as they do in wild type (Fig. 8A,B, green arrowheads).

Quantification of the distance from the hub to the DJ-GFP signal at the distal end of spermatid cysts revealed a significant increase in *isep^2-83^* mutants (Fig. 8C). Upon closer inspection, *isep^2-83^* sperm tails appeared broken and disorganized (Fig. 8A’’, B’’), resembling *orb2* null mutant (*orb2^36^*) sperm tails, which appear rough and irregular when stained for the protein Bol (Xu et al., 2012).

Polyglycylated tubulin staining, which marks axonemes from the onset of individualization (Bre et al., 1996, Bressac et al., 1995), returned similar results: axonemes did not extend as far back towards the hub (Fig. 7A, B, green arrowheads) and appeared broken and disorganized at the distal ends of the cysts, unlike in wild type (Fig. 7A,Bii-iii). When quantified, the hub-to-distal sperm tail distance was again significantly increased in *isep^2-83^* (Fig. 8D). These results indicate defects in axoneme elongation, in addition to defects in Soti localization.

Finally, we examined the RNA-binding proteins Orb2 and Orb, which mark the distal ends of cysts during and after elongation, respectively (Xu et al., 2012). In wild-type, Orb2 was most abundant at the distal ends of elongating cysts, which extend almost all the way to the hub (Fig. 8E, yellow and magenta arrowheads). In *isep^2-83^*, the distal ends did not extend as far toward the hub, and the Orb2 signal was less sharp (Fig. 8F, yellow and magenta arrowheads). Orb is detected near the hub once cysts have completed elongation but have not yet begun to individualize (Xu et al., 2012), accumulating strongly at the distal end of the cyst in a gradient similar to Soti (Fig. 8G, magenta arrowheads), In *isep^2-83^* the Orb signal was much further down the testis relative to the hub and was no longer sharp, adopting a “flared paintbrush” pattern like that of *orb2^ΔZZ^* mutants (Fig. 8H, yellow and magenta arrowheads) (Low et al., 2026).

Thus, using markers of actively elongating spermatids (Orb2), post-elongating but pre-individualization spermatids (Soti and Orb), and individualizing spermatids (DJ-GFP and polyglycylated tubulin), we observed that *isep^2-83^* cysts consistently failed to extend as far towards the hub as in wild type, and that Orb2 localization and the Orb and Soti gradients were less sharp. This suggests a role for *isep* in elongating spermatids and in establishing the Orb2, Orb and Soti gradients and raises the possibility that the individualization defects are secondary to defects in elongation. Consistent with this, Isep protein was co-expressed with Orb2 during late elongation, and with Orb post-elongation, but was not detected in individualizing cysts marked by DJ-GFP (Fig. 4B,D).

### TAIL Iso-seq reveals that transcripts with longer poly(A) tails in *isep^2-83^* mutants are enriched in GO terms for axonemal assembly, structure, and function

Since *isep* encodes a poly(A) polymerase, we asked whether the sterility defects correspond to changes in poly(A) tail length in the testis transcriptome. We isolated total RNA from four replicates each of *w^1118^* and of *w^1118^; isep^2-83^* mutant testes and performed TAIL Iso-sequencing (see Methods). The analysis yielded 17,189 unique transcripts across replicates and genotypes, including 1,421 novel transcripts (Table S1), and principal components analysis showed robust clustering of biological replicates by genotype (Fig. S3A). Rather than a decrease in global poly(A) tail lengths, as might be expected from loss of a poly(A) polymerase, we found a small but significant increase of 3 adenines in the median global tail length in *w^1118^; isep^2-83^* (Fig. 9A; Table S2).

**Figure 9:**
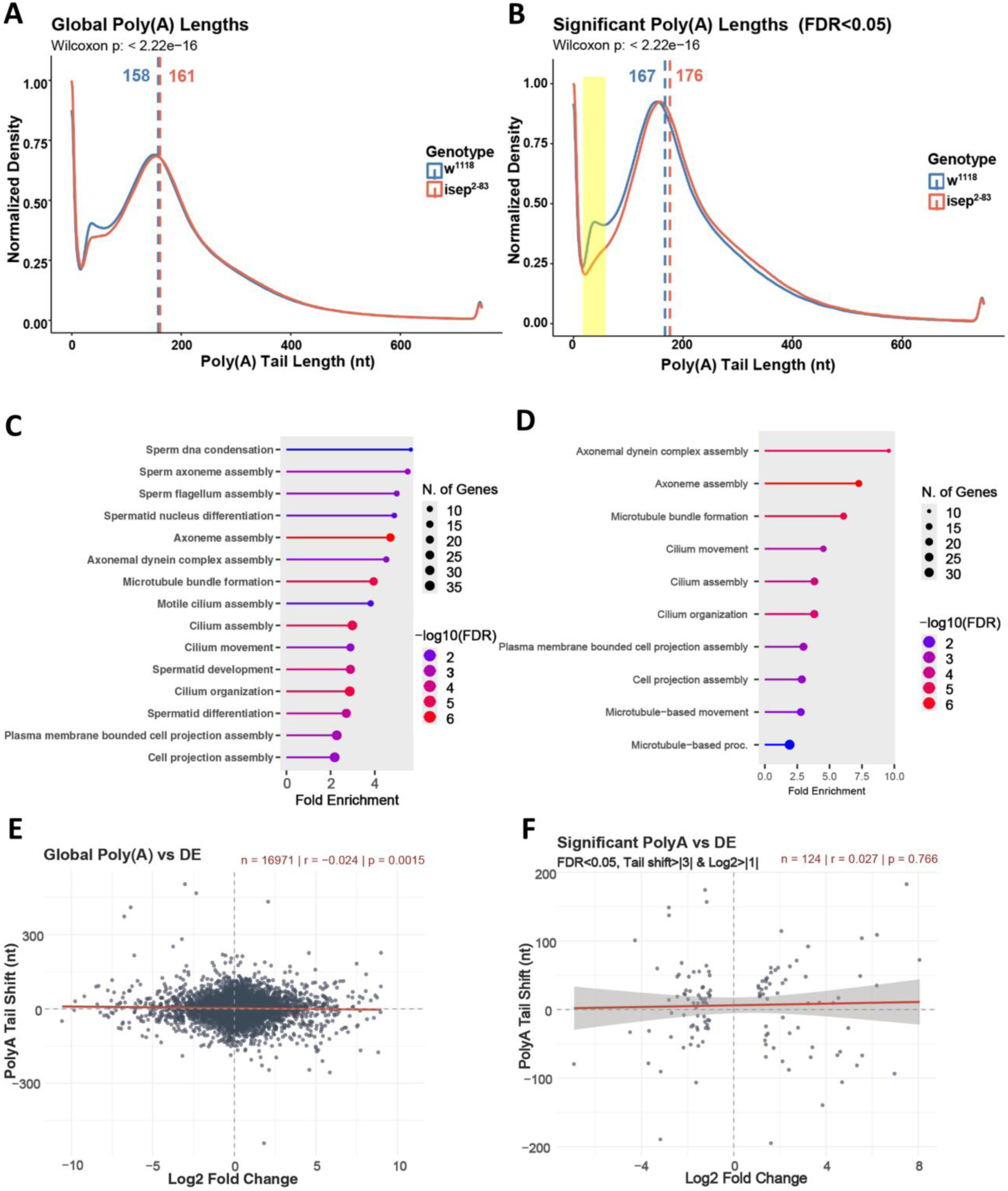
Tail Iso-seq reveals lengthened poly(A) tails in *isep^2-83^*mutants, enriched for spermatogenesis development and axonemal assembly and uncorrelated to changes in transcript levels. **(A)** Distribution of poly(A) tail lengths for all reads sequenced, and **(B)** for reads belonging to the significantly differentially polyadenylated (DPA) transcripts (tail shift >|3| adenines, FDR<0.05); yellow shading marks the 20-80 adenine population. **(C)** GO analysis of DPA transcripts with longer poly(A) tails in *w^1118^; isep^2-83^* vs. *w^1118^*and **(D)** those specifically lengthened above 80 adenines. **(E)** Poly(A) tail difference vs. log_2_ fold change for all detected transcripts and **(F)** for transcripts shared between the DPA and significantly differentially expressed data set. Number of transcripts, correlation coefficients, and P values are shown in the upper right.

We refined this global list to differentially polyadenylated (DPA) transcripts; those differing by at least 3 adenines and an FDR < 0.05, yielding 1,235 transcripts, 49 of them novel (Table S3). The distribution of reads for these transcripts shows a significant median increase of 9 adenines in *w^1118^; isep^2-83^*(Fig. 9B), and the majority were lengthened (960) rather than shortened (275). Isep therefore does not appear to be generally required to lengthen poly(A) tails in the testes but may be required to prevent hyperadenylation.

GO analysis of the full DPA dataset returned sperm axoneme assembly as the most enriched term, along with additional axoneme assembly terms and terms for spermatid development, sperm individualization, spermatid differentiation, spermatogenesis, and male gamete generation (Fig. S3B, Table S8). The DPA data set includes 29 of the 162 previously identified post-meiotically transcribed genes (Raz et al., 2023), including *isep-RC*. These GO terms align well with our phenotypic analysis, suggesting that Isep influences the polyadenylation of transcripts responsible for axoneme assembly.

Restricting GO analysis to transcripts with increased median tail length, the highest enriched term is sperm DNA condensation, with the remaining terms related to axoneme assembly and spermatid development (Fig. 9C, Table S8). This group includes the *Mst84Da-d* genes, which are required for normal axonemal structure and are known to change poly(A) tail lengths over spermatogenesis (Kuhn et al., 1991), and *isep-RC*, which could indicate that Isep regulates its own poly(A) tail length.

Conversely, transcripts with decreased median tail length returned one term: cytoplasmic translation (Fig. S3C, Table S8). Most of these encode ribosomal subunits, but the list also includes two eukaryotic translation initiation factors, *eIF4A and eIF3m*. The latter, also known as *Tango7*, encodes a protein that works with the apoptosome to remodel the syncytium during spermatid individualization; loss of *Tango7* results in male sterility and individualization failure (D’Brot et al., 2013). Thus, transcripts important for spermatogenesis are both hyper- and hypo-adenylated in *isep^2-83^*, suggesting that Isep’s effects on polyadenylation are complex.

The poly(A) length distribution of *w^1118^* shows the highest density of reads with a poly(A) tail length of 167 adenines, with another smaller peak of approximately 20-80 adenines, which is reduced in *w^1118^*; *isep^2-83^* (Fig. 9B, shaded area; Fig. S3D). We asked whether transcripts with reads in this range shifted towards shorter or longer tails in *w^1118^; isep^2-83^.* We identified all DPA transcripts with *w^1118^* reads in this range, binned the corresponding reads as <20, 20-80, and >80, and compared the percentage of each transcript’s reads falling into each bin (Table S4, see methods). Transcripts that had reads shifted significantly below 20 adenines or were stable showed no GO enrichment. Transcripts with reads significantly shifted above 80 adenines were enriched for terms related to axonemal and microtubule organization (Fig. 9D, Table S8) and include *Mst84Db-RA* and *isep-RC.* A distinct subset of 20-80 adenine transcripts was therefore lengthened in *w^1118^; isep^2-83^* and enriched for axonemal components, consistent with the axonemal defects we observe and with a requirement for *isep* in restricting hyperadenylation.

*TENT5* orthologs in mice and *C. elegans* preferentially polyadenylate transcripts containing ER signal peptides (Mackiewicz et al., 2026, Brouze et al., 2024, Mroczek et al., 2017). To determine whether this holds for *isep,* we retrieved amino acid sequences for the DPA transcripts (excluding novel transcripts and non-coding RNAs) and used SignalP 6.0 (Teufel et al., 2022). Signal peptides were predicted for 97 transcripts (9%), compared to the genome-wide rate of 22% (Duan et al., 2025). Of these, most had lengthened tails (68, 70%) rather than shortened (29, 30%) (Table S10), with no clear connections to spermatogenesis, indicating they are present by chance and that, unlike its ortholog, Isep’s mRNA targets are not ER-associated.

TAIL Iso-seq also reports transcript levels. Of the 17,189 transcripts, 17,144 passed DEseq2 filtering (Table S5). Globally, differences in transcript levels and poly(A) tail length showed a very weak correlation (Fig. 9E). DEseq2 identified 1,968 significantly differentially expressed transcripts (log_2_FoldChange >|1|, padj < 0.05), 566 of them novel (Table S6), with no significantly enriched GO terms. Of these, only 124 were also in the DPA dataset (Table S8); these too showed no correlation between changes in tail length and differential expression (Fig. 9F) and no GO enrichment. Thus, differences in polyadenylation observed in *isep* mutants do not appear to affect steady-state transcript levels, leaving open the possibility that Isep modulates transcripts by other post-transcriptional means, such as translational efficiency or localization.

Post-transcriptional regulation via polyadenylation relies on motifs primarily within the 3’ UTR and on the co-factors that bind to them. We therefore used MEME (Bailey and Elkan, 1994) and AME (McLeay and Bailey, 2010) to analyze the 3’ UTRs of the 1,134 DPA transcripts in *w^1118^; isep^2-83^*, and of 400 transcripts shifted above 80 adenines (excluding novel genes and non-coding RNAs lacking annotated UTRs). No prominent novel motifs were found in either data set. AME identified three known motifs in the DPA data set: binding sites for (1) Smooth (Sm), an ortholog of hnRNPL implicated in pre-mRNA processing, mRNA stabilization, and translation (Geuens et al., 2016) in 177 transcripts; (2) Cnot4, a sometime subunit of the CCR4-NOT deadenylase complex (Caulier et al., 2025) in 180, and (3) the SR protein B52, which affects polyadenylation site selection (Bradley et al., 2015) in 130 (Table S9). A fourth motif, inferred from human SC35 (Tacke and Manley, 1995), was also enriched but was not analyzed further. The list of DPA transcripts shifted above 80 adenines was enriched only for the Sm motif. Sm is particularly interesting, as loss-of-function alleles of *sm* cause male sterility (zur Lage et al., 1997); Sm localizes to the region where Y-loops are transcribed in spermatocytes, just like Isep (Lowe et al., 2014); and Sm is required for guidance of neuronal axons, which, like spermatid axonemes, are microtubule-based structures (Layalle et al., 2005). Given the overlap between Isep and Orb2 expression, we also scanned for Orb2 motifs, which we found in 91 DPA transcripts, including *isep-RC* (Table S9).

Additionally, we examined 5′UTRs. The four *Mst84D* genes and *Mst87F*, all present in the DPA data set, share a 12-base pair motif in their 5′UTRs that regulates their post-meiotic translation and cytoplasmic polyadenylation (Kuhn et al., 1991, Schafer et al., 1990). We find this motif in the 5’ UTRs of 79 DPA transcripts (Table S10). GO analysis returned terms for spermatid differentiation and development, but the most enriched term was again sperm axoneme assembly, with most remaining terms relating to axonemes and microtubules (Table S8), reinforcing the idea that *isep* regulates transcripts required for axoneme assembly.

Finally, analyzing whether these predicted motifs co-occur (Table S10) revealed a 5-fold enrichment of transcripts containing both a predicted Sm and Cnot4 motif, with 129 shared out of 177 and 180, respectively (Fig. S3E, Table S10). Thus, Isep, Cnot4, and Sm may work in concert to regulate specific transcripts. GO analysis of these shared transcripts yielded only one significant term, microtubule-based process (Table S9), and the list includes notable axonemal transcripts *Sak, Efa6,* and *Rsph3* (Qu et al., 2019, Bettencourt-Dias et al., 2005, Wang et al., 2019). Interestingly, *isep-RC* also contains predicted binding sites for Cnot4 and Sm.

Taken together, our transcriptomic analysis suggests that loss of *isep* leads not to a global shortening of poly(A) tails but to lengthening. The dysregulation does not correlate with transcript levels, and transcripts with lengthened tails are enriched for genes involved in axonemal assembly and other microtubule-based processes.

## DISCUSSION

*isep (CG46385)* encodes the sole *Drosophila melanogaster* ortholog of the *TENT5* family of ncPAPs. Although we initially identified *isep* as a potential transcriptional target of the Pax6 ortholog, Eyeless, *isep* mutants have superficially normal eyes. Instead, *isep* is essential in the male germline, where its loss causes male sterility: spermatids fail to individualize, and actin cones are no longer ‘in step’ with each other.

### Isep is required for elongation and individualization

The individualization failure in *isep^2-83^* homozygotes is unambiguous: mutant cysts show uniform cleaved caspase rather than restriction to the cystic bulge; the bulges are elongated and flattened; waste bags are absent; nuclei are scattered; and actin cones scatter at the onset of individualization and fail to travel synchronously down the cyst.

Isep also appears required for proper axoneme elongation. Isep is expressed at the distal ends of cysts during elongation, before individualization begins. Mutant spermatid tails appear broken and disorganized 1) during active elongation when they express Orb2; 2) at the end of elongation when they express Orb and Soti; and 3) after individualization when they express DJ-GFP and polyglycylated tubulin. Consistent with this, transcripts with elongated poly(A) tails in *isep* mutants are enriched in GO terms for axonemal assembly and spermatid development.

Isep may be required separately for both elongation and individualization. However, the individualization failure may be a downstream consequence of elongation defects. Because elongation establishes the cyst’s cellular architecture, mutations in genes active during this phase typically cause individualization failures (Texada et al., 2008, Xu et al., 2024, Fabrizio et al., 1998).

### Isep resembles post-transcriptional gene regulators in spermatogenesis

Isep’s expression pattern and mutant phenotypes suggest links to factors involved in post-transcriptional gene regulation during spermatogenesis, including the CPEB orthologs Orb and Orb2 and the testis-specific mRNA cap-binding protein eIF4E-5. All are expressed at the distal ends of cysts, with Isep colocalizing with Orb2 during active elongation (Xu et al., 2012), and with Orb in the gap between active elongation and individualization (Xu et al., 2012, Kaplan et al., 2010), while eIF4E-5 appears just before and during individualization (Shao et al., 2023).

The phenotypes are correspondingly similar. Both *orb2* and *eIF4E-5* are required for individualization and elongation (Shao et al., 2023, Kaplan et al., 2010, Xu et al., 2012, Low et al., 2026). *orb2*, *eIF4E-5,* and *isep* are required for proper Soti localization at distal cyst ends, and *orb2* and *isep* are required for proper Orb localization (Low et al., 2026). In *orb2* mutants, the distal cyst ends have the same “flared paintbrush” appearance (Low et al., 2026) as in *isep* mutants (Figure 8H), a phenotype attributed to disrupted cyst organization and polarization that leads to mislocalization of Orb and Soti (Low et al., 2026). Thus, Isep may participate with Orb2, Orb, and eIF4E-5 to regulate post-transcriptional gene expression late in spermatogenesis. The similarity to *orb2* is particularly striking, and *isep-RC* contains a predicted consensus CPEB-binding site. Since both *orb* and *soti* are known Orb2 targets, Orb2 may regulate all three factors.

Isep’s role appears distinct from that of the other ncPAP required for spermatogenesis, Gld2. Although both localize to the distal ends of cysts, *Gld2*-knockdown testes show basal body docking defects, actin cones never form, and protamine incorporation is disrupted (Sartain et al., 2011). Moreover, candidate Gld2 targets that had no changes in poly(A) tails in *Gld2* knockdowns appear in our DPA dataset, with *Mst77F, Mst87F, chic, jar,* and *yuri* all having longer poly(A) tails, while only *bol* was shorter.

### Does Isep function as a poly(A) polymerase during spermatogenesis?

Like other *TENT5* members, Isep retains the conserved nucleotidyltransferase motifs essential for poly(A) polymerase activity (Hu et al., 2020), and vertebrate TENT5s show this activity *in vitro* (Hu et al., 2020, Mroczek et al., 2017, Zhang et al., 2021). However, our data argue that polyadenylation is unlikely to be Isep’s primary biochemical function in spermatogenesis. First, although some transcripts have shorter tails in *isep* mutants, and some of these have known roles in spermatogenesis, the majority have longer tails. Second, a straightforward model in which Isep polyadenylates targets to promote their stability is not supported, as changes in poly(A) tail length did not correlate with changes in transcript levels. Third, a predicted catalytically dead Isep rescued the sterility phenotype to the same extent as wild type. In contrast, TENT5C appears to require poly(A) polymerase activity for its function in spermatogenesis in mice (Baptissart et al., 2026), although apparently not for all its roles (Kazazian et al., 2020).

Whatever Isep’s biochemical role, Isep clearly influences poly(A) tail length during spermatogenesis, both positively and negatively, on transcripts enriched for roles in axonemal elongation and spermatogenesis, which fits our phenotypic evidence. As with *isep*, whole-testis poly(A) sequencing in mice showed that TENT5C/D loss also dysregulated poly(A) tails, predominantly toward longer tails, but with tail shortening of particular transcripts (Brouze et al., 2024). In contrast, more recent profiling of a specific isolated germ cell population in mice found that global poly(A) distributions of round and elongated spermatid transcriptomes were almost identical between wild-type and TENT5C mutants (Baptissart et al., 2026), with significant poly(A) tail shortening in a specific set of transcripts. Whether the discrepancy in global poly(A) tail length between whole testes and specific germ cell populations results from population-specific tail lengths or loss of specific cell populations from TENT5 mutant testes is unclear. Nevertheless, although Isep does not appear to require poly(A) polymerase activity for its role in spermatogenesis, whereas mouse TENT5C does, Isep seems to share with its vertebrate counterpart a role in regulating poly(A) tail length.

### Poly(A) tail shortening is associated with active translation

The longer poly(A) tails observed in both *isep* mutants and mouse *TentC/D* mutants may result from hyperadenylation of mRNAs. However, recent results in tissue culture cells show that deadenylation follows ongoing translation rather than preceding it (Shu et al., 2025), and short tails are a conserved feature of highly expressed genes (Lima et al., 2017). Thus, the longer poly(A) tails in *isep* mutants could result from a failure to initiate translation of stored transcripts rather than from being hyperadenylated. The same is true in mice, where, in round spermatids, stored transcripts encoding late spermatogenesis factors such as protamines and transition proteins have long poly(A) tails that shorten as they are translated during elongation (Baptissart et al., 2026), and in *Drosophila* where deadenylation of Protamine A and B correlates with translation initiation (Sartain et al., 2011).

### Isep may organize a translational hub at the distal end of spermatid cysts

If Isep doesn’t function as a poly(A) polymerase, how might it affect poly(A) tail length? Our DPA transcripts are enriched for predicted 3’ UTR binding sites for Sm, B52, and Cnot4. Some also carry predicted Orb2 binding sites and a 5’ UTR motif shared by the *Mst84D* cluster and *Mst87F*, and some of these sites co-occur significantly. This suggests that Isep might interact with transcripts via these RNA-binding proteins, even though it lacks an RNA-binding domain (Liudkovska and Dziembowski, 2021).

We propose that Isep functions less as an enzyme acting on individual transcripts but rather as a factor required for proper assembly and positioning of a complex within which transcripts involved in spermatogenesis are regulated. Isep has intrinsically disordered regions (IDRs) at both termini, and its C-terminal glutamine-rich region is prion-like. Low-affinity interactions between such regions are thought to drive phase separation and assembly of biomolecular condensates (Gomes and Shorter, 2019). These condensates could include Orb2, Cnot4, and Sm, all of which possess IDRs with polyglutamine regions, along with the CCR4-NOT deadenylase complex, GLD2/TENT2, and the two CPEBs. Because such a complex could contain both deadenylases and ncPAPs, its disruption could account for transcripts with both shorter and longer poly(A) tails. Our observed co-localization of Isep with the CPEBs Orb2 and Orb supports this idea.

Accordingly, translational regulation by the CPEB2-4 subfamily, which includes Orb2, involves phase separation mediated by their N-terminal IDRs. In their unphosphorylated state, these proteins partition into microtubule-associated condensates that sequester and repress bound mRNAs, whereas hyperphosphorylation disassembles these condensates and promotes translation (Fernandez and Mendez, 2025). Furthermore, Orb associates with P-granules (Kanke et al., 2015), and drives polyadenylation-dependent local translation of pericentrin-like protein at centriole-derived structures in *Drosophila* embryos (Fang and Lerit, 2022).

Microtubules themselves can serve as a platform for condensate formation (Volkov and Akhmanova, 2024). Growing spermatid axonemes could thus provide the physical scaffold that nucleates and anchors this complex, where *isep, soti, orb,* and other cup and comet transcripts accumulate, acting as a microtubule-associated hub in which stored, long-tailed transcripts interact with translation machinery. Loss of *isep* would then impair translation of axonemal transcripts, producing the phenotypes we observe. Interestingly, vertebrate TENT5C also associates with microtubule-based structures, both in roles that do not require its poly(A) polymerase activity (Chen et al., 2020, Kazazian et al., 2020), and in its roles in spermatogenesis (Zheng et al., 2019, Baptissart et al., 2026). Moreover, in both mouse *Tent5c* mutants and *Drosophila isep* mutants, proteins fail to localize properly to microtubule-based structures, with mouse ODF1 failing to concentrate at the head-tail linkage (Baptissart et al., 2026) and Soti and Orb failing to accumulate at the distal ends of axonemal microtubules in *isep^2-83^*mutants. This suggests a conserved role in spatial control of translation, although the substrates and landmarks diverge.

## DATA AND REAGENT AVAILABILITY

Gene expression data, fly strains, and plasmids generated in this study are available upon request.

## ACKNOWLEDGMENTS

We thank Dr. Samyi Belteton, Director of Microscopic Imaging at the NMSU RESEARCH CORES PROGRAM, for his expert assistance with the imaging techniques used in this study. We also extend our gratitude to Dr. Nasser Rusan (Cell and Developmental Biology Center, National Heart Lung and Blood Institute, National Institutes of Health) for generously providing the topi-Gal4 line, and to Dr. Eli Arama (Professor at the Weizmann Institute of Science) for providing the Anti-Scotti antibody. *Drosophila* microinjections were performed by Rainbow Transgenics. Finally, we acknowledge the Developmental Studies Hybridoma Bank (DSHB) for providing critical antibodies used in this work and the Bloomington Drosophila Stock Center for flies.

## AI STATEMENT

AI was used to improve the language of the manuscript Claude (Anthropic) and for generating and debugging code to analyze the TAIL-seq data Gemini (Google). All outputs were reviewed and accepted, modified, or rejected, and all code was tested and its outputs verified.

## MATERIALS AND METHODS

### Oligos and Vectors

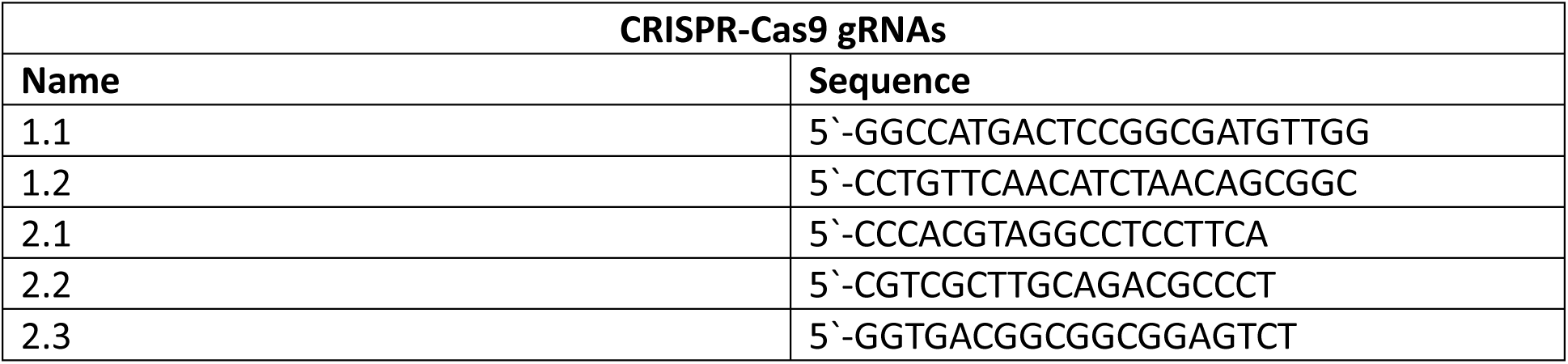
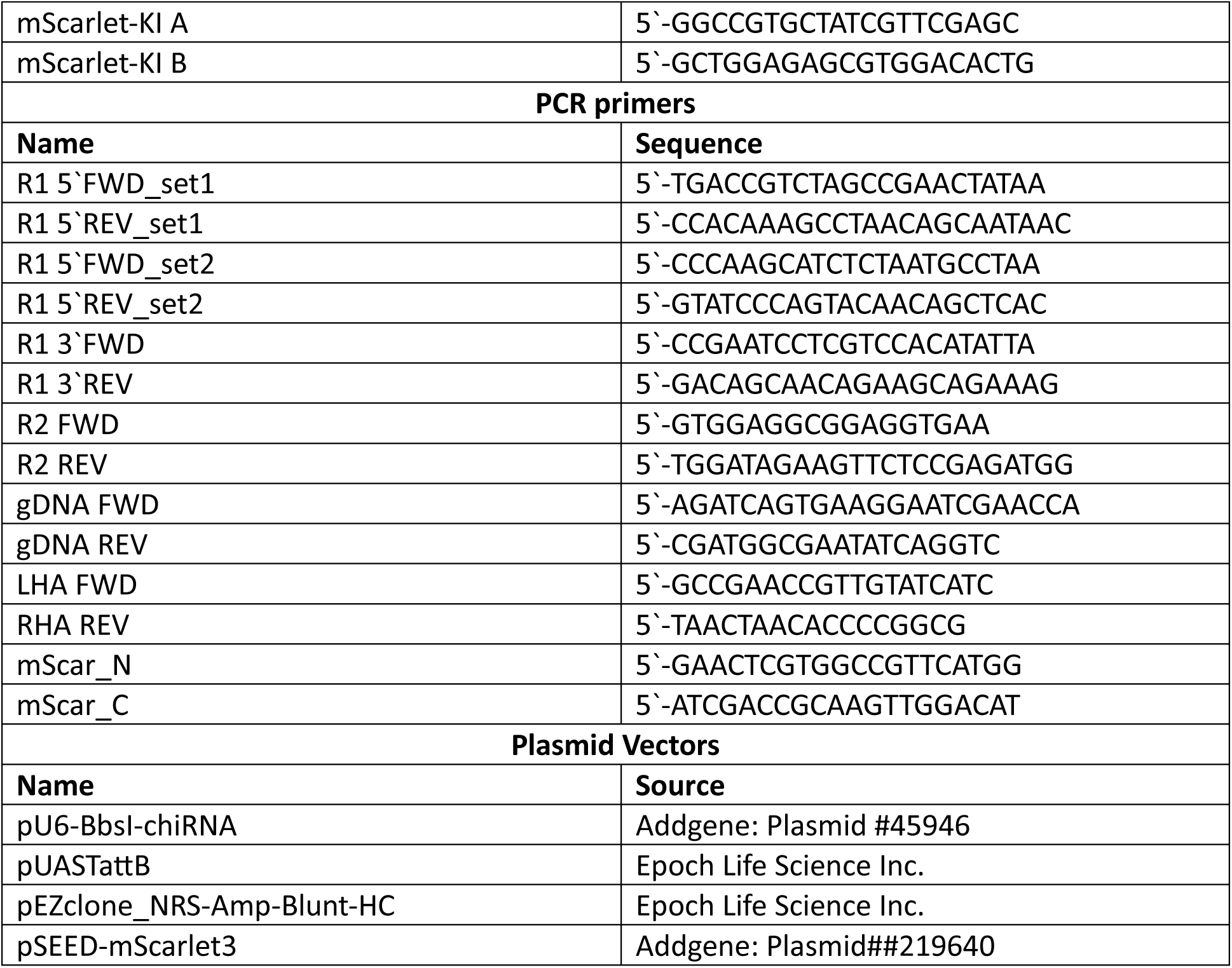

### Fly Strains

- Wild-type control: *Oregon R*
- w[1118]
- *Mi[MIC]CG46385^MI01603^*; genomic coordinates: 2R:7,746,162..7,746,162 [-] (Venken et al., 2011, Nagarkar-Jaiswal et al., 2015).
- *Mi[MIC]CG46385^MI03850^*; genomic coordinates: 2R:7,765,950..7,765,950 [+] (Venken et al., 2011, Nagarkar-Jaiswal et al., 2015).
- *Mi[MIC]CG46385^MI12561^*; genomic coordinates: 2R:7,781,696..7,781,696 [+] (Venken et al., 2011, Nagarkar-Jaiswal et al., 2015).
- w[1118]; Df(2R)Exel6054, P{w[+mC]=XP-U}Exel6054/CyO (BDSC: 7536, Parks et al., 2004)
- w[1118]; Df(2R)ED1725, P{w[+mW.Scer\FRT.hs3]=3’.RS5+3.3’}ED1725/SM6a (BDSC: 8941, Ryder et al., 2007)
- w[*]; P{w[+mC]=protamineB-eGFP}2/CyO; P{w[+mC]=dj-GFP.S}3/TM3, Sb[1] (BDCS: 58406, Xiangyi Lu, Wayne State University)
- w[1118]; Dp(2;3)GV-CH321-77L09, PBac{GV-CH321-77L09}VK00031/TM3, Sb[1] (BDCS: 90598, GenetiVision Corporation, 2016-)
- y, sc, v; +/+; nos-Cas9 (Rainbow Transgenic Flies Inc: nos-Cas9 attP2)
- M{vas-int.Dm}ZH-2A, P{CaryP}attP2 (Rainbow Transgenic Flies Inc: R8622)
- Topi-Gal4 (kindly gifted from Nasser Rusan) (Raychaudhuri et al., 2012)
- Asl-GFP (kindly gifted from Nasser Rusan) (Galletta et al., 2020)
- (Harvester) w; Dr[1]/TM3, Sb[1] Ser[1] P{w[+mC]=attR.nos-Cas9, U6:3-1+2sgRNA.attRS18 (BDCS: 604565, Aguilar et al., 2024)

### CRISPR-Cas9 mutagenesis

CRISPR-Cas9 mutagenesis for generation of the three *isep* alleles was performed in two rounds. In the first round, two guide RNAs (1.1 and 1.2) were injected simultaneously. In round two, three guide RNAs (2.1-2.3) were injected simultaneously. Generation of gRNA expression plasmids was via the U6-gRNA (chiRNA) cloning protocol outlined in https://flycrispr.org/. Rainbow Transgenic Flies performed the injections into *nos-Cas9 attP2* flies. Sequences of guide RNAs used for CRISPR-Cas9 mutagenesis are listed in Oligos and Vectors. We screened resulting mutations by PCR using primer sets R1 (round 1) and R2 (round 2) (Oligos and Vectors), then Sanger sequenced them using the same primer sets.

*CG46385^1-54^* was generated in CRISPR-Cas9 round 1, and is a 281-nucleotide deletion (2R: 7,778,276--7,778,556) that removes the splice site at the end of the first coding exon of isoforms RA and RE with a one-base-pair insertion. This is predicted to change codon 303 in isoforms PA and PE, resulting in a stop codon six codons later.

*CG46385^2-1^* is a 27-nucleotide deletion (2R: 7,741,342--7,741,368) that removes amino acids RRIYTCSLK (codons 515-523 in isoforms RA and RE, codons 198-206 in isoforms RB and RF, and codons 167-175 in isoforms RC and RD).

*CG46385^2-83^* is a 70-nucleotide deletion (2R:7,741,343--7,741,412) with a five-nucleotide insertion (CCCAC). This causes a frameshift (at codon 500 for isoforms PA and PE, codon 183 for isoforms PB and PF, and codon 152 for isoforms PC and PD) that results in a stop codon 17 codons later.

### Isep-mScarlet CRISPR/Cas9 knock-in

mScarlet-3 was knocked into the endogenous *isep* gene between the N-terminal intrinsically disordered region and the beginning of the folded region (between amino acids T393 and Q394 based on isoform PE). In brief, we performed a knock-in using the SEED/Harvest method (Aguilar et al., 2024). Homology arms (below) flanking the insertion sites and containing silent mutations (lowercase letters) in the gRNA sites were synthesized *de novo* by Epoch Life Sciences and then cloned into the pEZ vector (Oligos and Vectors). We modified the mScarlet3 vector to replace the eGFP screening marker with the *miniwhite* gene from the pUASTattb vector; we used Golden Gate Assembly (NEB, #E1601) to insert *mScarlet3/miniwhite* into the pEZ vector containing the homology arms. mScarlet-KI A and mScarlet-Kl-B guide RNAs were cloned into the pU6-BbsI-chiRNA vector as described above. Flies positive for the screening marker *miniwhite* were crossed to flies containing the germline harvester as described in (Aguilar et al., 2024) and screened for correct harvesting by the absence of *miniwhite* followed by PCR screening using primers gDNA FWD, gDNA REV, LHA, RHA, mScar_N, and mScar_C (Oligos and Vectors). Correct harvesting was confirmed by Sanger sequencing using these same primers.

Left homology arm:

CACTAGACGGAGCCGAACCGTTGTATCATCAACAGCGCAGCAACAGAATGGACGTCTACCAGATCGACGATGGATTCCACTCGGACGGCGGCACCGAGACTCCGCCGCCGTCACCCAGTTCTGGGTCATCGATCGCCTCGACCTCGGACCACGGATCcttgGAatccGTGGAtACcGGcGGGACG

Right homology arm:

CAGCGGCTGGCgGTatTGAGtTTtGAaCAaGTGAGCAAACTGCACGACGTCATGGACGAGAAGGTGGCCATCCATGGGCGCGGCAATTTTCCCACACTGGAGGTGAAGCTCAAGGATCTGGTCAACCTGGTGCGCCGCAAGCTGGAGGCGGAGGTGAACGCCGGCGGCGCCGGGGTGTTAGTTAA

### Egg lay assay

0–3 day-old males and virgins of the desired genotype were crossed to *w^1118^* males and virgins, and *w^1118^*males and virgins were crossed to each other as the control. Each cross consisted of three males and five females. Egg-laying chambers consisted of 50ml conical tubes with holes made in the sides for ventilation. The caps of the tubes were filled with an agar/molasses mixture (40% molasses, 0.05% agar) with a dollop of yeast paste (baker’s yeast plus water mixed to a creamy consistency) in the middle. Flies were placed in the chamber and allowed to lay eggs for 24 hours, after which they were removed from the chambers. We counted the number of eggs laid, then counted the number of eggs hatched at 12 and 24 hours after the adults were removed. Each cross consisted of three replicates.

### Live squash preparation of the testis

Testes were dissected in testis isolation buffer (TIB; 183 mM KCl, 47 mM NaCl, 10mM Tris pH 6.8) with Hoechst 33342 tri-hydrochloride tri-hydrate (Life Technologies H3570) (1:1000) and transferred onto poly-L-lysine-coated slides with a drop of TIB and Hoechst. We cut off the tip of each testis and placed a coverslip on top. Cells were drawn out of the sheath by removing excess TIB from around the coverslip with a Kimwipe. Images were acquired on an inverted Leica DMi8 epifluorescence microscope with a 40x phase-contrast objective and Leica K5 camera using the Thunder Imaging System.

### HCR fluorescent *in situ* hybridization

A 1463 bp sequence corresponding to the catalytic domain of *Isep* (2R: 7,741,822..7,740,360) was submitted to Molecular Instruments, Inc. to generate 20 split-initiator probe pairs, and *wingless* probes were purchased commercially from Molecular Instruments, Inc. Third instar larval tissue and adult testes from 0–3 day-old males were dissected in 1X PBS, then fixed (4% PFA, 1X PBS, 0.1% Triton-X100) for 30 minutes at room temperature, rinsed 3 times with PBT (1X PBS, 0.1% Tween-20), and then washed 3 times for 15 minutes with PBT. For larval tissue, the remainder of the procedure was performed according to the manufacturer’s instructions for generic samples in solution (https://files.molecularinstruments.com/MI-Protocol-RNAFISH-GenericSolution-Rev9.pdf).

For adult testes, Hoechst was added after amplification of hairpins (step 5 in the protocol). After amplification, the testes were washed once for 5 minutes with 5X sodium citrate buffer with Tween 20 (SSCT) (0.75 M sodium chloride, 75mM sodium citrate, 0.1% Tween 20), followed by a 15-minute wash with probe wash buffer. Hoechst 33342 tri-hydrochloride tri-hydrate (Life Technologies H3570) was diluted 1:800 in probe wash buffer and incubated with the testes for 2 hours at room temperature.

Testes were then washed once for 15 minutes with probe wash buffer, followed by one 5-minute wash, two 30-minute washes, and another 5-minute wash with 5X SSCT. Samples were imaged using a Leica TCS SP5 II confocal microscope.

### Immunofluorescence

Testes from 0–3 day-old males were dissected in 1X PBS and fixed (4% PFA, 1X PBS, 0.1% Triton X-100) for 30 minutes at room temperature. Samples were permeabilized (0.3% sodium deoxycholate, 1X PBS, 0.3% Triton X-100) twice for 15 minutes, then washed once with PBT (1X PBS, 0.5% Triton X-100) for 10 minutes. Samples were then blocked (3% BSA, 1X PBS, 0.1% Triton X-100) for 30 minutes at room temperature before being incubated overnight at 4°C in the blocking solution with the primary antibody. Samples were then washed three times for 5 minutes with PBT and incubated with secondary antibodies for 2 hours in the dark at room temperature in blocking solution. Samples were then washed three times for 5 minutes with PBT before post-fixation (2% PFA, 1x PBS, 0.1% Triton X-100). Samples were washed three more times for 5 minutes with PBT before being mounted in 80% glycerol with 4% n-propyl gallate. Primary antibodies used were mouse monoclonal anti-Orb 1:10 (DSHB, 6H4), mouse monoclonal anti-Orb2 1:10 (DSHB, 4G8), rabbit polyclonal anti-cleaved caspase-3 1:400 (cell signaling #9661s), guinea pig polyclonal affinity –purified anti-Soti 1:100 (kind gift of Eli Arama, Department of Molecular Genetics, Weizmann Institute of Science), mouse monoclonal anti-pan polyglycylated Tubulin Antibody, clone AXO 49 1:500 (Sigma-Aldrich, MABS276), and rabbit polyclonal anti-RFP 1:400 (Rockland, 600-401-379). Secondary antibodies and stains used were Alexa Fluor™ 568 Phalloidin 1:400 (Invitrogen A12380), Hoechst 33342 tri-hydrochloride tri-hydrate 1:800 (Life Technologies H3570), with all secondaries used at 1:200; anti-mouse Alexa Fluor™ 647 (Invitrogen A21237), anti-mouse Alexa Fluor™ 488 (Jackson ImmunoResearch code:715-545-150), anti-guinea pig Cy5 (Jackson ImmunoResearch code:706-175-148), anti-rabbit Alexa Fluor™ 488 (Jackson ImmunoResearch code: 111-545-003), and anti-rabbit Alexa Fluor™ 647 (Jackson ImmunoResearch code: 111-605-003). Samples were imaged using a Leica TCS SP5 II confocal microscope.

### Rescue using the Gal4-UAS system

Two UAS lines were generated: the first, a codon-optimized coding sequence (CDS) of isoforms RC/RD, *UAS-isep^WT,^* and the second, the same except with a missense mutation of aspartic acid 130 (in isoform RC/RD) to alanine, as our poly(A) catalytic dead, *UAS-isep^CD^*. Epoch Life Sciences Inc. synthesized the CDSs *de novo* and subcloned them into the pUASTattb vector, which contains the screening marker mini-white. The resulting plasmids were subsequently injected into embryos expressing PhiC31 integrase under the control of the *vasa* promoter and harboring an attP site on the third chromosome (Rainbow Transgenic Flies Inc., line: R8622). We obtained males that were homozygous for *isep^2-83^* and contained one copy of either *UAS-isep^WT^* or *UAS-isep^CD^*, and *topi-Gal4.* As controls, we used *isep^2-83^* with only Gal4 as a negative control, and flies with either UAS construct and Gal4 but lacking the *isep^2-83^* allele as positive controls. 30 single male crosses with two *w^1118^* females were set up for each genotype. We recorded the time to the first pupa, then counted pupae after three days; if pupae appeared, the cross was considered fertile. All crosses were allowed to go for 30 days before the experiment was terminated.

### Testis RNA isolation

For each replicate, 120 testes from 3-5-day-old flies were dissected in cold 1X PBS and immediately placed into TRI Reagent® (Sigma-Aldrich, 93289), homogenized, and stored at −80°C for up to 1 week until all samples were completed. 0.2 volumes of chloroform were used to isolate the aqueous layer, which was combined with 1.5 volumes of 100% ethanol. RNA from the aqueous/ethanol mixture was then isolated using the MagMAX™ mirVana™ Total RNA Isolation Kit (ThermoFisher, A27828), loaded onto the KingFisher™ Apex automated purification system, and run using the “High-throughput isolation of RNA (including small RNA) from serum and plasma samples”, skipping the proteinase K digestion.

### Tail Iso-Seq profiling and analysis

A full-length cDNA library was generated by CD Genomics using the PCR-cDNA Barcoding Kit (SQK-PCB111.24, Oxford Nanopore Technologies) and sequenced using the Nanopore platform. Raw fast5 binary data were generated containing all original sequencing signals, where each read corresponds to a single fast5 file. After base calling by Dorado, fast5 files were converted to fastq files. Pychopper (2.2.2) (https://github.com/nanoporetech/pychopper) was used to identify full-length reads with a quality score threshold of 7 and a minimum length of 50 bases. NanoFilt (2.8.0) (De Coster et al., 2018) was subsequently applied to filter these reads further, ensuring an average quality score of 7 or higher and a minimum length of 50 bases. Minimap2 (2.24-r1122) (Li, 2018) was used to align the cleaned reads to the *Drosophila melanogaster* genome (version r6.67). Salmon (1.9.0) (Patro et al., 2017) was used to estimate read counts, map each alignment to a unique isoform, and assign an FBtr ID. If no confident isoform-level mapping could be performed, then the alignment was assigned to its parent gene and classified using a FBgn ID. FlyBase ID counts were then filtered for all downstream analysis to include only those transcripts that had a count per million of 0.5 or greater in at least 3 out of the 4 biological replicates for either *w^1118^* or *w^1118^; isep^2-83^*. DEseq2 (1.34.0) (Love et al., 2014) was used for differential gene expression. For the poly(A) dataset, we filtered out all reads that were −1, which indicates a failed poly(A) length estimate. This data was then compiled by genotype, and for each transcript, the number of reads remaining, median, and mean poly(A) tail length were calculated. We compared median tail lengths of *w^1118^; isep^2-83^* to *w^1118^* using a two-tailed Mann-Whitney U test (Wilcoxon rank-sum test) for significance, and calculated FDR using the Benjamini-Hochberg test.

For the shift analysis, we identified all DPA transcripts (FBtr or FBgn IDs) with *w^1118^* reads between 20 and 80 adenines and collected all reads for those transcripts from both *w^1118^* and *w^1118^*; *isep^2-83^*. We analyzed only transcripts with at least 20 reads in each genotype. We binned reads for each transcript as 0-19, 20-80, or 81+ adenines, and calculated the percentage of a transcript’s reads in each bin for each genotype. We compared read counts per bin between genotypes using a Pearson’s chi-squared test of independence, and calculated FDR using the Benjamini-Hochberg method. We calculated the change in each bin as the percentage in *w^1118^*; *isep^2-83^* minus the percentage in *w^1118^*. Because transcripts with many reads can reach significance with small differences, transcripts with FDR < 0.05 were called stable if neither the 0-19 nor the 81 and above bin gained more than 5 percentage points and the 20-80 bin lost no more than 5. We called the remaining transcripts with FDR < 0.05 shifted up or shifted down based on whether the 81 and above or the 0-19 bin gained the larger share of reads in *w^1118^*; *isep^2-83^*. Transcripts with no reads in the 0-19 bin in either genotype could not be tested and are marked as not tested in Table S5.

We performed all GO analyses using ShinyGO 0.85 (Ge et al., 2020), using the filtered counts per million dataset as the background. We used the MEME suite to scan for novel motifs (Bailey and Elkan, 1994). We used Analysis of Motif Enrichment (AME) with the *Drosophila melanogaster* CISBP-RNA (Catalog of Inferred Sequence Binding Preferences of RNA-binding proteins) database (McLeay and Bailey, 2010) to scan for motifs of known RNA-binding proteins. We used Find Individual Motif Occurrences (FIMO) to scan for the Orb2 and Mst87F motifs (Grant et al., 2011). We predicted signal peptides using the SignalP 6.0 prediction tool (Teufel et al., 2022).

## SUPPLEMENTAL FIGURE

**Supplemental Figure 1:**
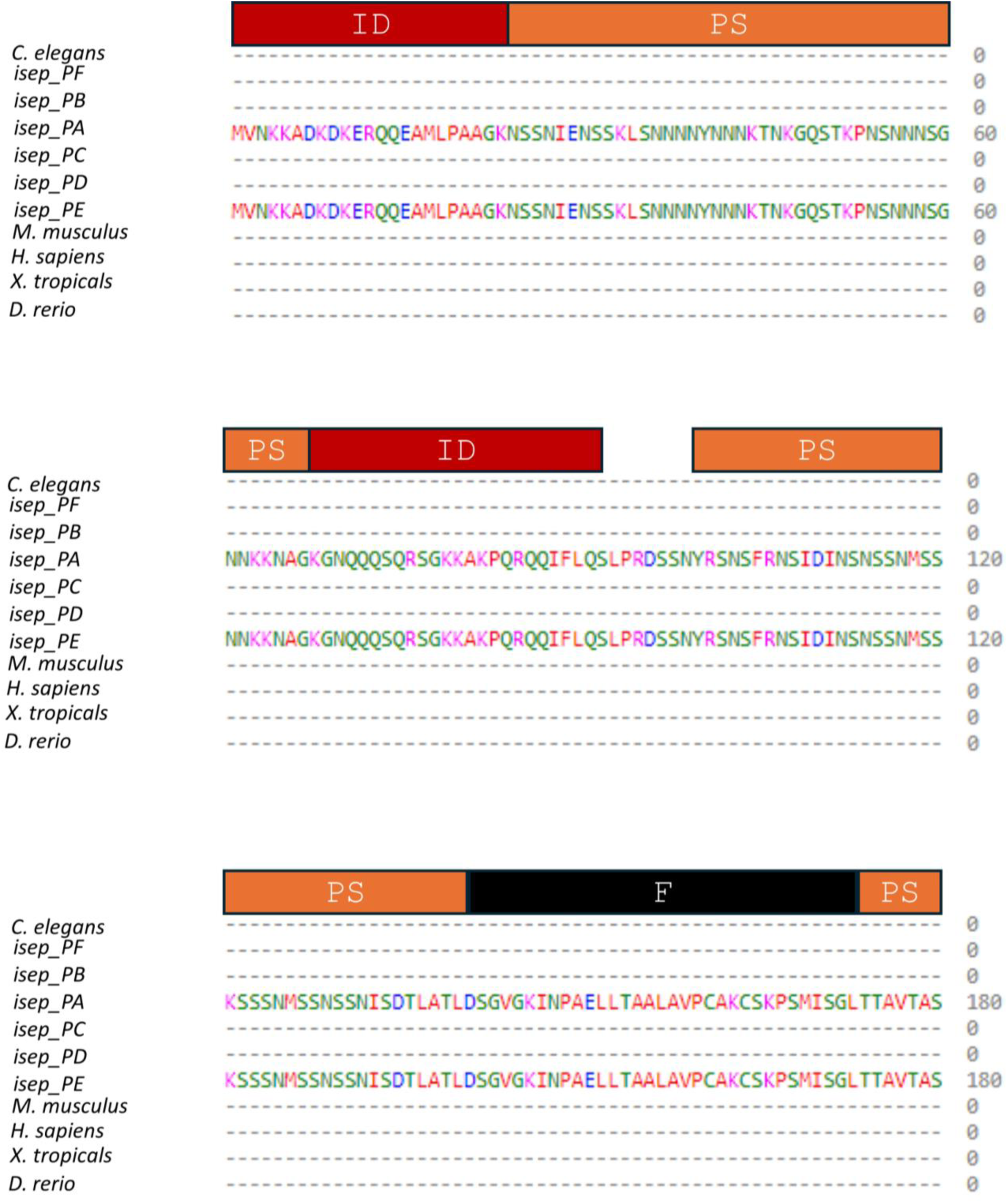
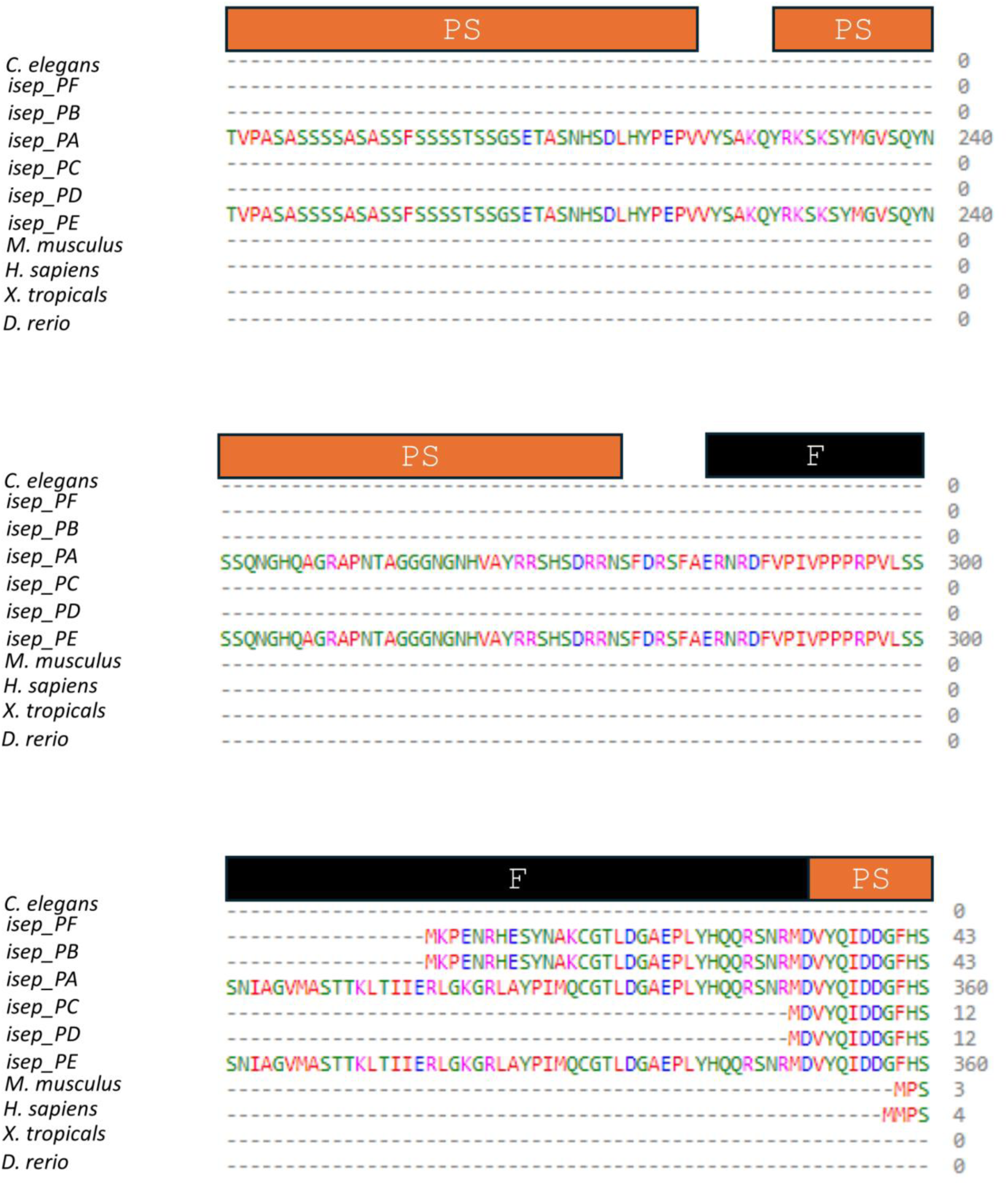
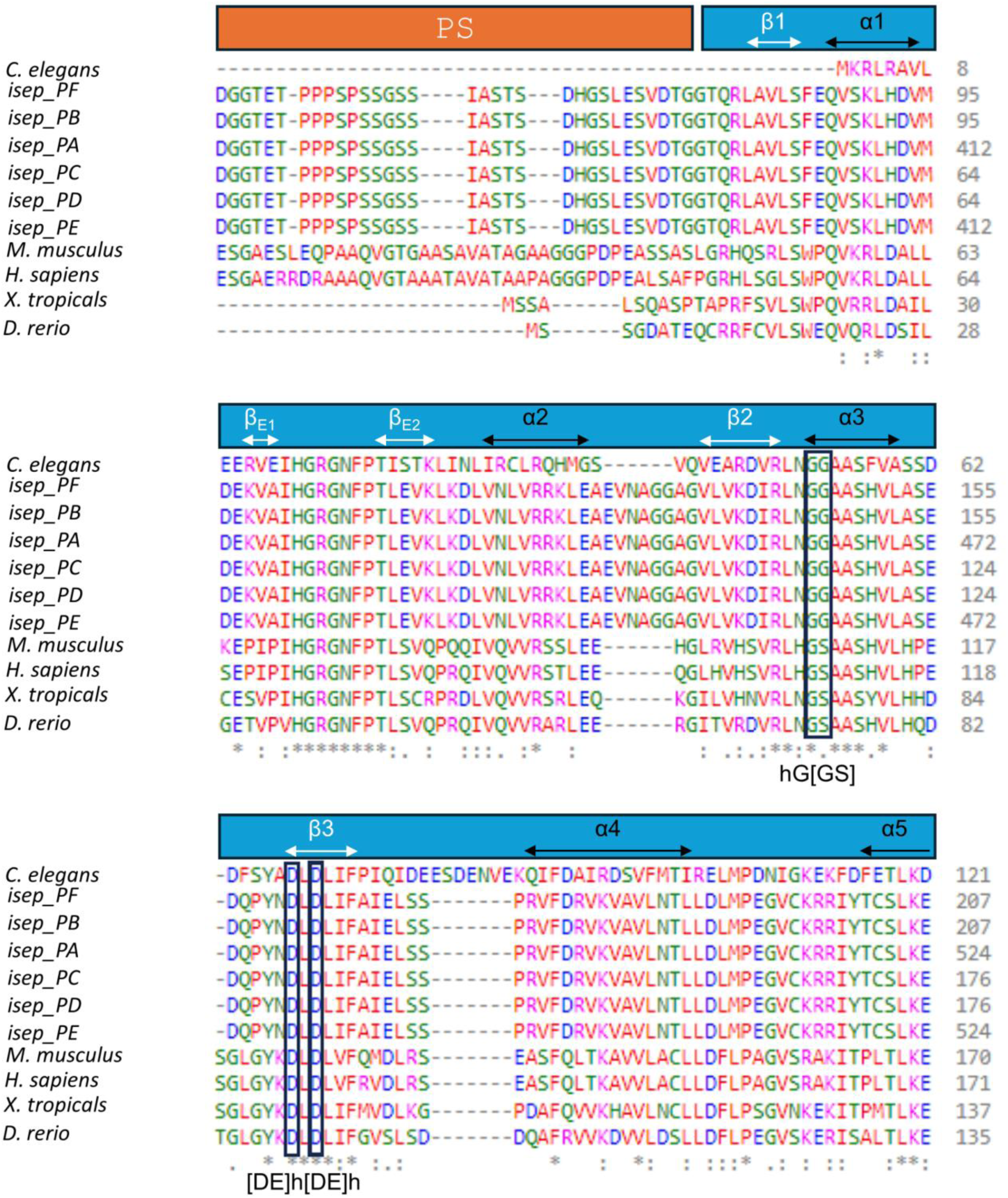
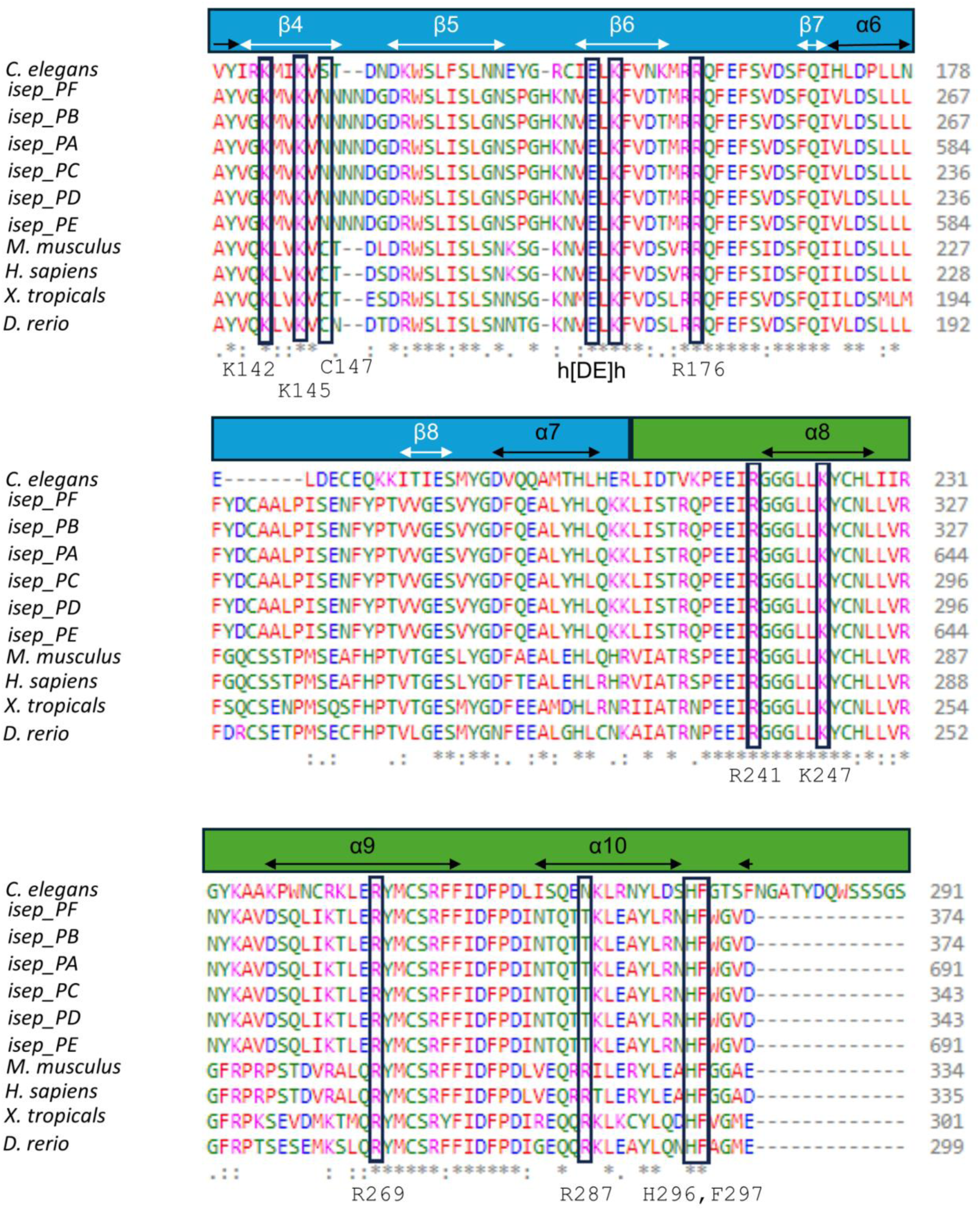
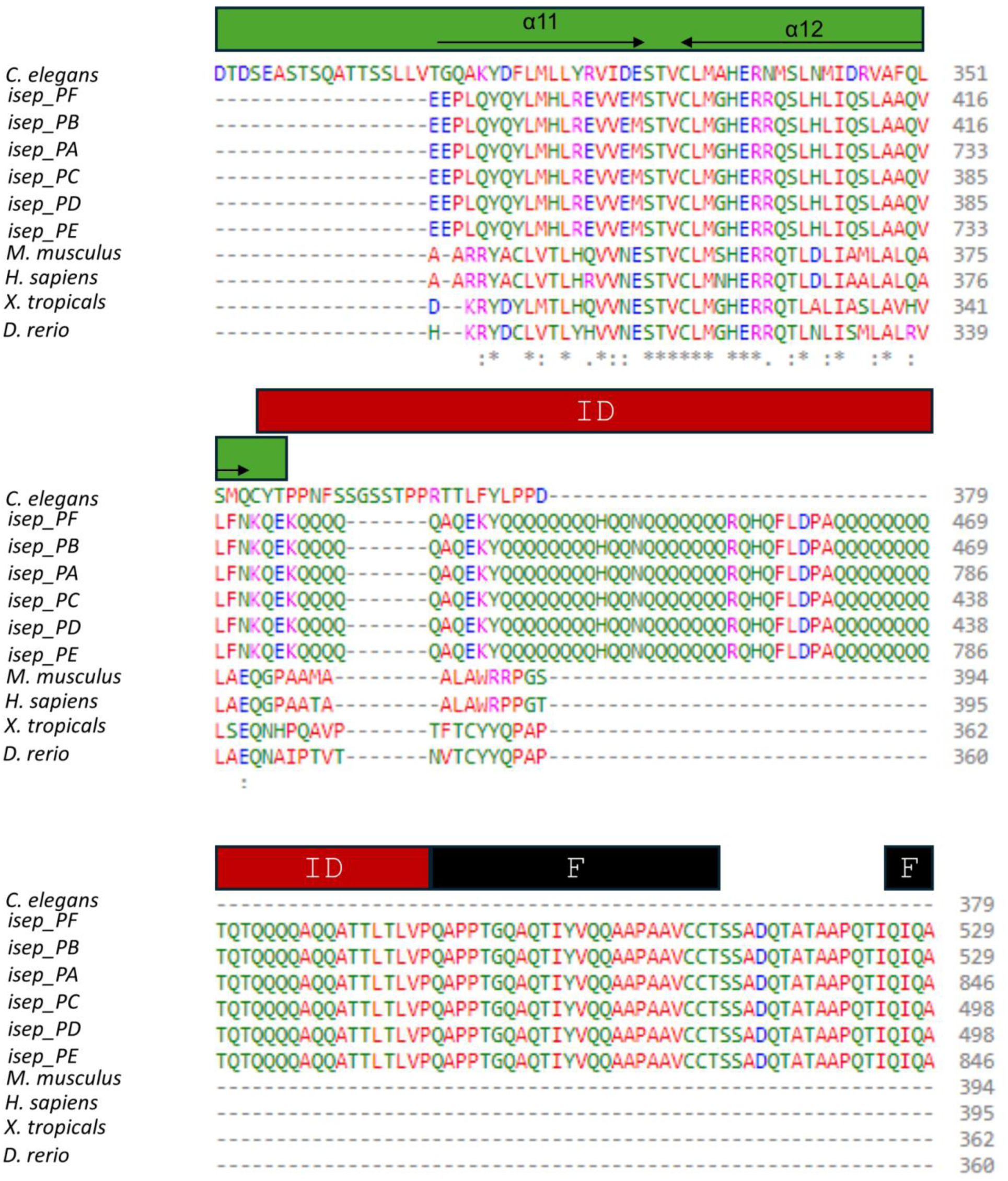
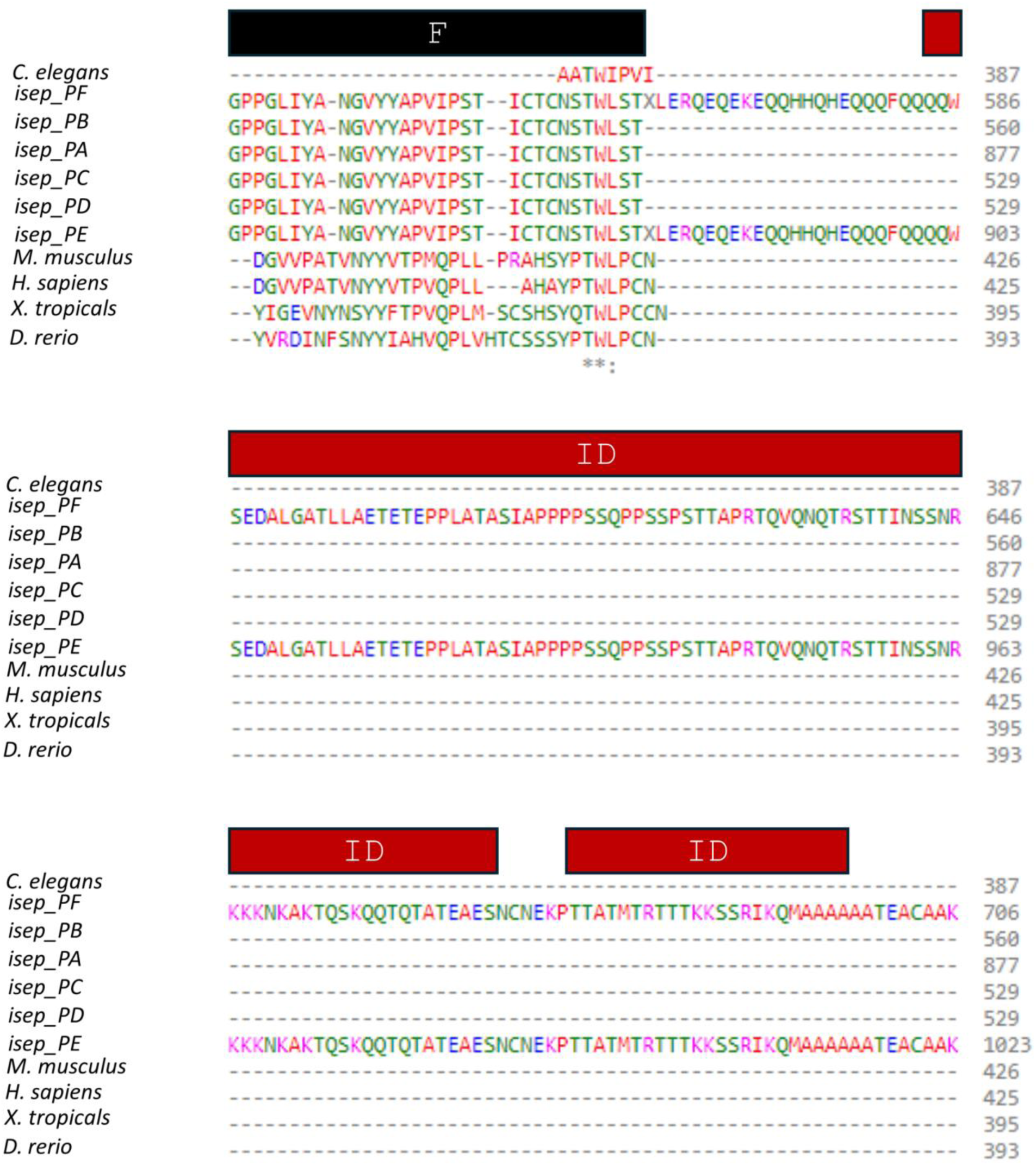
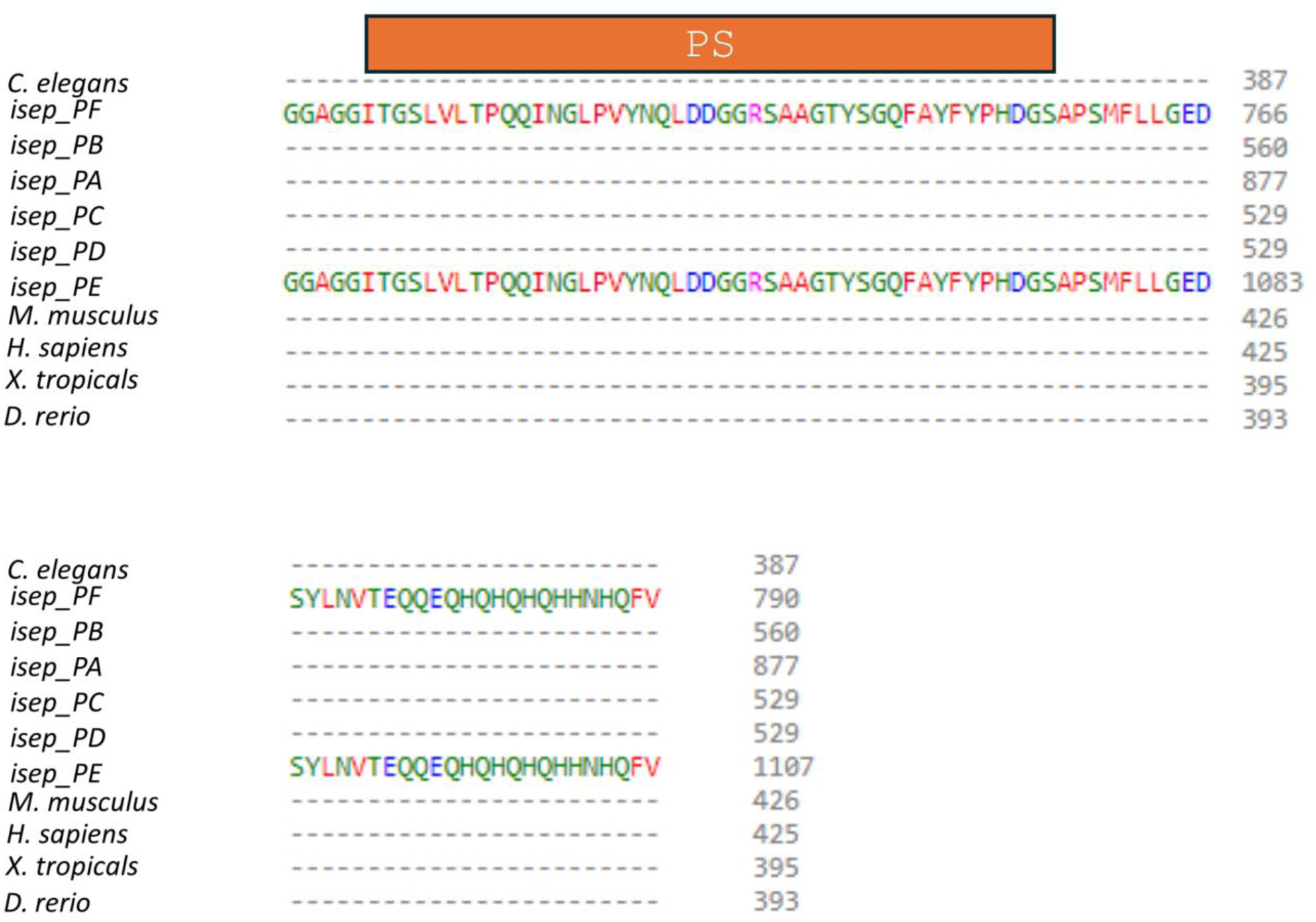
*isep* encodes a TENT5 ortholog. Clustal Omega alignment of *D. melanogaster* predicted protein isoforms (Isep PA-PE) with *Caenorhabditis elegans* (*C. elegans*) TENT-5 and with TENT5B from *Mus musculus* (*M. musculus*), *Homo sapiens* (*H. sapiens*), *Xenopus tropicalis* (*X. tropicalis*), and *Danio rerio* (*D. rerio*). Red bars above the alignments mark regions of Isep predicted by ParSe 2.0 (Wilson et al., 2023) to be intrinsically disordered; orange bars mark predicted intrinsically disordered regions that can also undergo phase separation; and black bars mark predicted folded regions. Blue and green bars above the sequence indicate secondary structure elements for the N-terminal catalytic domain (NCD) (light blue) and the C-terminal helical domain (HD) (green) from a crystal structure of *X. tropicalis* TENT5B (Hu et al., 2020), which together form a cleft in the tertiary structure. Black boxes mark the five residues required for TENT5B’s poly(A) polymerase activity (Gly107, Ser108, Asp124, Asp126, Glu200); alanine substitutions abolish poly(A) polymerase activity (Hu et al., 2020). All five are conserved in Isep. Of the eight positively charged residues implicated in TENT5B RNA binding, seven are conserved in Isep (K142, K145, K169, R176, R241, K247, H296) and one (R287) is not. Residues whose substitution has little or no effect on TENT5B activity are less well conserved: R269 and F297 are retained in Isep, whereas C147 is not. The first intrinsically disordered region on the C-terminal side of the HD is glutamine-rich, and PANTHER (Thomas et al., 2022) accordingly classifies Isep as a prion-like Q/N-rich domain-bearing protein, comparable to the C-terminal regions in vertebrate TENT5C. The *isep^2-1^* deletion (Figure 1C) is predicted to remove part of the linker between the fourth and fifth alpha helices of NCD together with the N-terminal portion of the fifth helix.

**Supplemental Figure 2:**
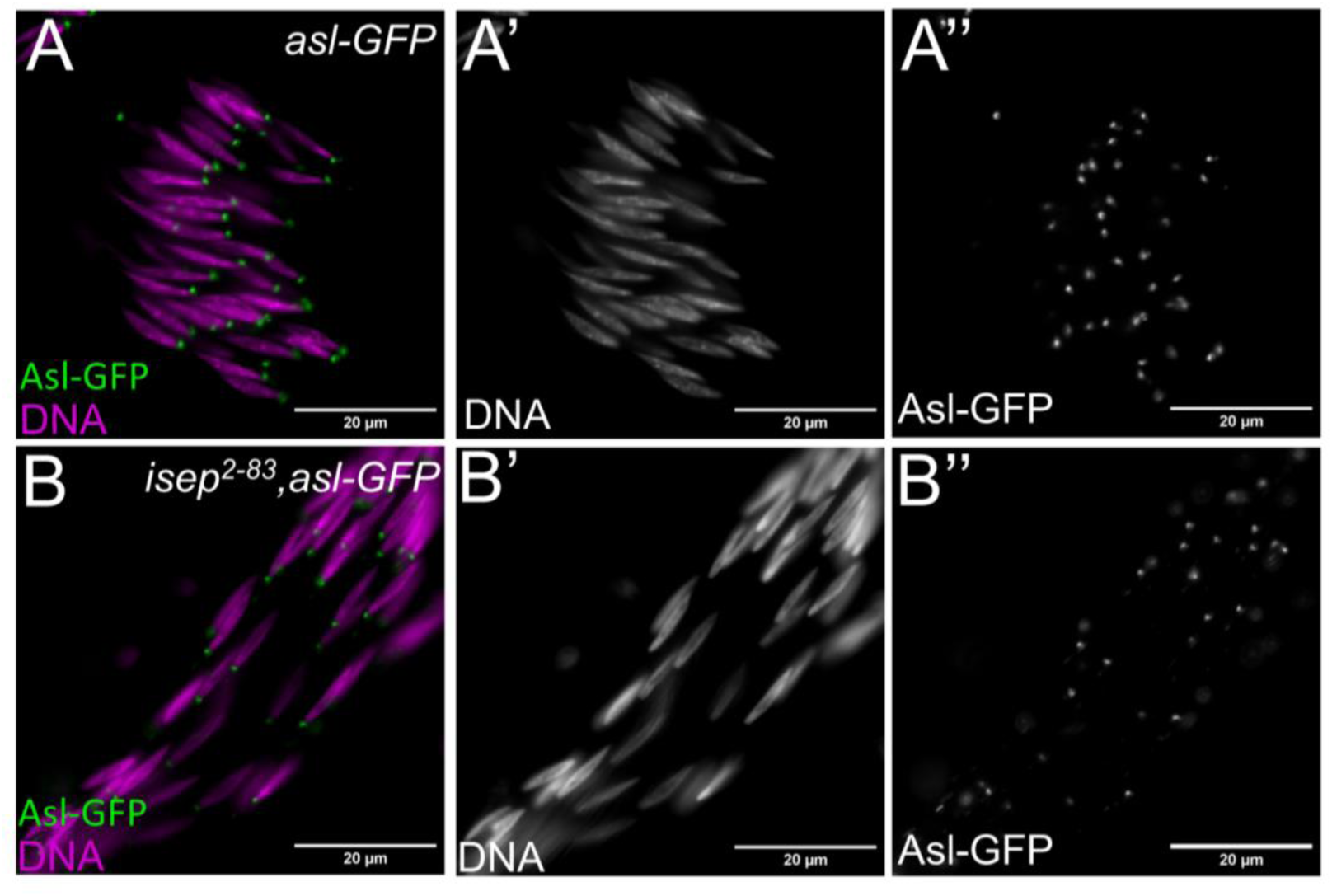
No obvious basal body defects were found in *isep^2-83^*mutants. Live squashed preparations of **(A)** *asl-GFP* (green)and (**B**) *isep^2-83^, asl-GFP* (green) spermatids stained for DNA (Hoechst, magenta) to visualize basal bodies and nuclei, respectively.

**Supplementary Figure 3:**
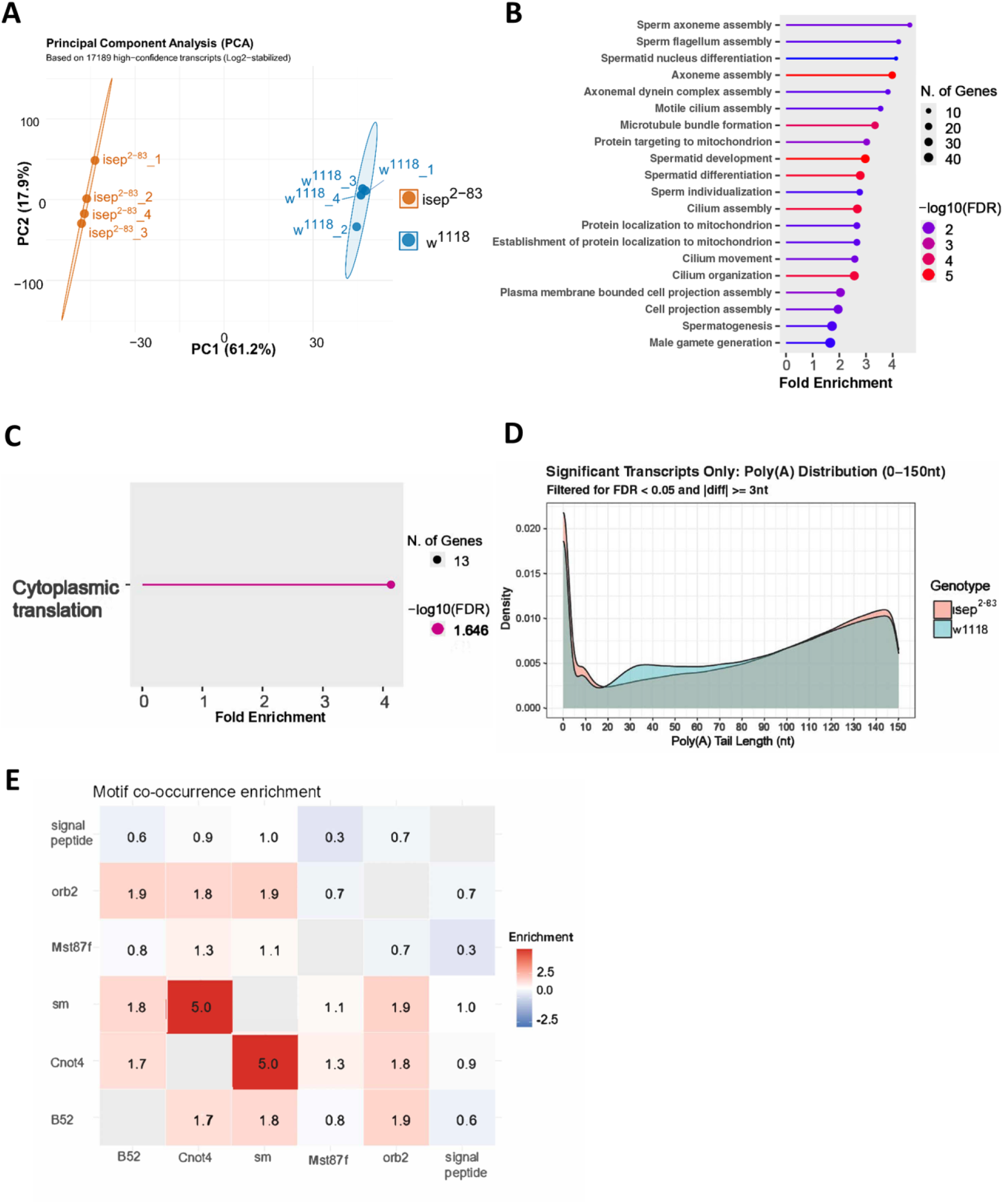
PCA and GO analysis of DPA list, with motif co-occurrences. **(A)** PCA of each biological replicate for *w^1118^; isep^2-83^* and *w^1118^.* **(B)** GO analysis of all transcripts in the DPA data set. **(C)** GO analysis of all transcripts in the DPA data set that had shorter poly(A) tails in *w^1118^; isep^2-83^ vs. w^1118^.* **(D)** Distribution of poly(A) tail lengths of individual reads *w^1118^; isep^2-83^*and *w^1118^* from 0-150 adenines. **(E)** Heat map showing enrichment of motif co-occurrences of transcripts in the DPA dataset.

## SUPPLEMENTARY TABLE LEGENDS

**Supplementary Table 1: Transcripts detected by TAIL Iso-seq.** All transcripts with a count per million of at least 0.5 in three or more of the four biological replicates in *w^1118^; isep^2-83^* or *w^1118^*.

**Supplementary Table 2: poly(A) tail measurements for all detected transcripts in *w^1118^; isep^2-83^* and *w^1118^.*** Median and mean poly(A) tail lengths for all detected transcripts.

**Supplementary Table 3: poly(A) tail measurements for differentially polyadenylated (DPA) transcripts.** Transcripts with a tail-length change of at least 3 adenines and an FDR < 0.05 in *w^1118^; isep^2-83^* and *w^1118^*.

**Supplementary Table 4: Shift analysis.** DPA transcripts with *w^1118^* reads between 20-80 adenines and the corresponding transcripts in *w^1118^*; *isep^2-83^*, with read counts and percentages in the <20, 20-80, and >80 adenine bins for each genotype (see Methods). Delta columns give the change in percentage (*w^1118^*; *isep^2-83^* minus *w^1118^*). Primary Defect: **Significant but Stable**, FDR < 0.05, but neither the <20 nor the >80 bin gained more than 5 percentage points and the 20-80 bin lost no more than 5; **Shifted Up** (>80 nt) or **Shifted Down** (<20 nt), all other transcripts with FDR < 0.05, according to whether the >80 or <20 bin gained the larger share of reads; **Not Significant**, FDR ≥ 0.05; **Not tested**, no reads below 20 adenines in either genotype. Because calls are based on which end bin gained reads, a transcript can be called shifted without losing reads from the 20-80 bin, and a transcript called stable can still show a shift of reads from the >80 bin into the 20-80 bin.

**Supplementary Table 5: DE-seq2 results of all transcripts analyzed.**

**Supplementary Table 6: DE-seq2 results of all transcripts that were significantly differentially expressed between *w^1118^; isep^2-83^*and *w^1118^***.

**Supplementary Table 7: Transcripts that were both significantly differentially polyadenylated and differentially expressed**. Further categorized based on shorter or longer tails and whether down- or up- regulated in *w^1118^; isep^2-83^* vs. *w^1118^*.

**Supplementary Table 8: All GO analyses performed; all GO terms with an FDR<0.05 are in bold. (Sheet: Whole DPA)** All transcripts in the DPA data set. **(Sheet: DPA up)** All transcripts that had longer poly(A) tails in *w^1118^; isep^2-83^* vs *w^1118^*. **(Sheet: DPA down)** All transcripts that had shorter poly(A) tails in *w^1118^; isep^2-83^* vs *w^1118^*. **(Sheet: Shifted up 80<)** All transcripts shifted by more than 80 adenines in *w^1118^; isep^2-83^* vs *w^1118^*. **(Sheet: Mst87F motif)** All transcripts found to contain a predicted 5′UTR motif identified in the *Mst87F* and *Mst84D* cluster. **(Sheet: Cnot4-Sm)** All transcripts that contain both a predicted Cnot4 and Sm binding motif in their 3′UTR.

**Supplementary Table 9: All predicted motifs found and predicted signal peptides**. **(Sheet: Signal peptide)** All transcripts in the DPA list were analyzed for predicted signal peptides, with ‘yes’ and ‘no’ denoting whether one was present, along with whether that transcript’s poly(A) tail was shorter (down) or longer (up) in *w^1118^; isep^2-83^* vs *w^1118,^* and what the actual length change was. Each transcript was grouped by parent gene, and the number of isoforms present in the DPA list is noted. **(Sheet: Orb2 3′ motif and Sheet: Mst87F 5′ motif)** FIMO results of DPA transcripts that contain a predicted Orb2 binding site or motif found in the 5′UTR of *Mst87F*, with the motif ID from the CISBP-RNA database for Orb2 and the position weight matrix used for both on the bottom. Each row is one motif occurrence, so a transcript can appear more than once, and transcripts assigned only at the gene level (FBgn) appear with several UTR sequences (suffixes _1, _2, and so on) that were counted as one transcript. **(Sheet: 3′motif search)** AME results for all 3′UTRs of transcripts from the DPA data set.

**Supplementary Table 10: Motif co-occurrence with predicted signal peptides. (Sheet: association stats)** Each motif binary combination is compared to the number of transcripts that contain that motif from the DPA data set, how many are shared, what percentage of transcripts containing that motif is also found with the one being compared against, the lift (ratio of observed to expected co-occurrence of the two motifs), the odds ratio of finding a transcript within the DPA data set that contains both motifs, and the p-value and adjusted p-value using a Fisher’s exact test. **(Sheet: gene list)** Summary table of all transcripts from the DPA list marked with a yes whether they contain any of the motifs analyzed.

